# Phloem loading via the abaxial bundle sheath cells in maize leaves

**DOI:** 10.1101/2020.09.06.284943

**Authors:** Margaret Bezrutczyk, Nora R. Zöllner, Colin P. S. Kruse, Thomas Hartwig, Tobias Lautwein, Karl Köhrer, Wolf B. Frommer, Ji-Yun Kim

## Abstract

Leaves are asymmetric, with differential functionalization of abaxial and adaxial tissues. The bundle sheath (BS) surrounding the vasculature of the C3 crop barley is dorsoventrally differentiated into three domains: adaxial structural, lateral S-type, and abaxial L-type. S-type cells seem to transfer assimilates towards the phloem. Here we used single-cell RNA sequencing to investigate BS differentiation in C4 maize. Abaxial BS (^ab^BS) cells of rank-2 intermediate veins specifically expressed three SWEET sucrose uniporters (*SWEET13a, b*, and *c*) and UmamiT amino acid efflux transporters. *SWEET13a, b, c* were also identified in the phloem parenchyma (PP). Thus maize acquired a unique mechanism for phloem loading in which ^ab^BS cells provide the main pathway for apoplasmic sucrose transfer towards the phloem. This pathway predominates in veins responsible for phloem loading (rank-2 intermediate), while rank-1 intermediate and major veins export sucrose from the phloem parenchyma (PP) adjacent to the sieve element companion cell (SE/CC) complex, as in Arabidopsis. We surmise that ^ab^BS identity is subject to dorsoventral patterning and has components of PP identity. These observations provide first insights into the unique transport-specific properties of ^ab^BS cells and support for a modification to the canonical phloem loading pathway of maize, which may be generalizable to other C4 monocots.

## INTRODUCTION

Leaves are typically asymmetric: there are often differences in the relative stomatal and trichome densities and cuticle properties between the abaxial and adaxial leaf surfaces. While maize leaves are amphistomatic, asymmetry remains apparent, with bulliform cells only on the adaxial side and a conjoint, collateral and closed-type vasculature, with adaxial xylem and abaxial phloem. Dorsoventral patterning in maize leaf is initiated in the shoot apical meristem at the earliest stages of leaf primordia development by expression of *RGD2* (*Ragged seedling2*) (Henderson et al., 2006) and adaxial expression of *RLD1* (*Rolled leaf1*), which is conferred by miRNA166-mediated *RLD1* transcript cleavage on the abaxial side (Juarez et al., 2004b, 2004a). This pattern is maintained throughout development by specific localization of numerous transcription factors, including the abaxial expression of *KANADI* (Candela et al., 2008). Bundle sheath cells (BSCs) of maize are not known to be functionally differentiated. In barley leaves, the BSCs are anatomically distinct: abaxial side “L-type” BSC have large chloroplasts, while “S-type” BSCs, with small chloroplasts, surround the rest of the mestome sheath. It was proposed that the S-type cells may be specialized for photoassimilate transport, based on the rapid disappearance of starch after the light period and the abundant plasmodesmatal connections between S-type cells, mestome sheath, and phloem (Williams et al., 1989). In maize, the two abaxial BSCs (^ab^BSCs) are typically smaller compared to the medial BSCs. *In situ* hybridization and immunolocalization had shown that Rubisco, the glutamine synthetase isoform GS1-4 (*GLN3*), and malic enzyme localized specifically to BS, with transcripts and proteins equally represented in all BSCs (Langdale et al., 1988a; Martin et al., 2006). Here we used single cell RNA sequencing (scRNA-seq) to test whether the BS of the C4 plant maize is uniform or also shows a dorsoventral differentiation of its cells, as found in barley. The analysis identified mesophyll- and bundle sheath-specific transcripts and found that the BS could be subclustered into two groups, one specifically expressing a variety of genes including SWEET13 sucrose transporters, UmamiT and AAP amino acid transporters, as well as several transcription factors. *In situ* hybridization and analysis of translational GUS fusions demonstrated that the subclustering was due to dorsoventral patterning, as evidenced by the finding that the three SWEET13 paralogs were specifically expressed in ^ab^BS. These findings not only show that the maize leaf BS is functionally differentiated, but also identify a unique pathway for apoplasmic phloem loading in a C4 plant. In addition, the three SWEET13s were also present in cells that most likely represent the phloem parenchyma, similar to the profiles of their Arabidopsis orthologs AtSWEET11, 12, and 13. Maize ^ab^BS thus appears to use a combination of dorsoventral patterning and PP identity to drive sucrose into the apoplasm of the phloem.

## RESULTS

### mRNA patterns of specific cell types in maize leaves

To determine whether maize leaves contain multiple BSC types, we performed single-cell sequencing on protoplasts isolated from maize leaves. We first established a protocol for protoplast release. To minimize the possibility of a developmental gradient across cells, fully differentiated tissue was harvested from the distal portion of leaf 2 of late V2 stage plants (first and second leaf collar exposed) (Li et al., 2010) (Fig. 1a). Standard leaf protoplasting protocols leave intact bundle sheath ‘strands,’ consisting of BSCs and the vasculature (Kanai and Edwards, 1973; Langdale et al., 1989). In a parallel study, we were able to optimize protoplasting of Arabidopsis leaves to increase the yield of vascular cell types (Kim et al., 2020). We compared published protocols and varied parameters such as incubation time, enzyme concentration, enzyme blend, and preincubation in L-cysteine (Ortiz-Ramírez et al., 2018). Efficiency of the release of putative vascular cells was monitored using qRT-PCR with cell-type specific markers, under the assumption that the *SWEET13* paralogs, analogous to their Arabidopsis orthologs, would be specific to phloem parenchyma. Although none of the protocols were able to yield efficient release of putative vascular cells, we obtained apparent release of both BSCs and vascular cells with Protocol 4 (Supplementary Fig. 1).

**Figure 1.**
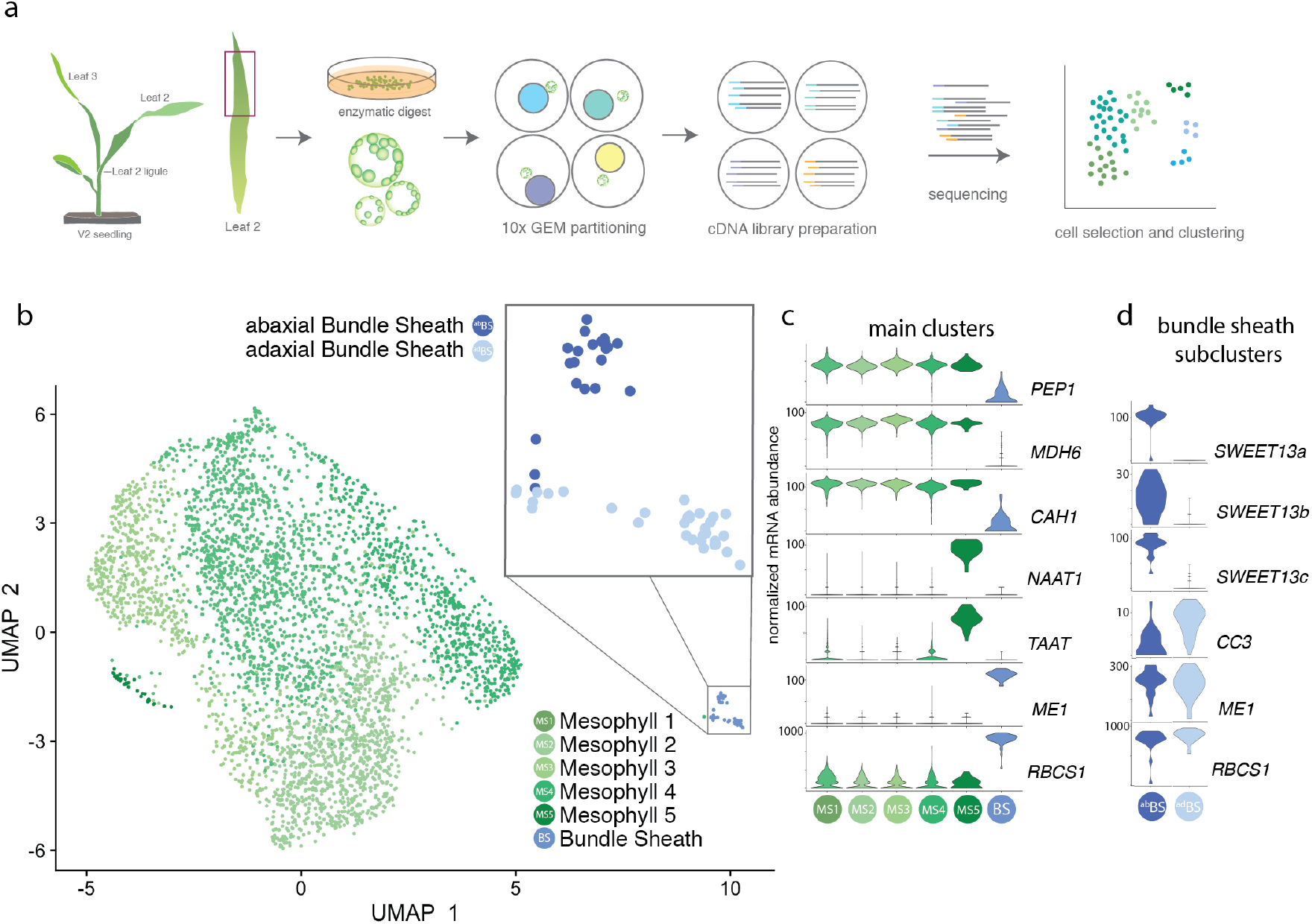
Mesophyll and bundle sheath clusters show canonical expression of C4 photosynthesis genes. **a.** Illustration of protoplasting, 10x Chromium Gel Bead-in-Emulsion (GEM) partitioning and cDNA synthesis, RNA sequencing, and data analysis. **b.** UMAP plot showing 2D representation of cell relationships in multidimensional space; bundle sheath cells separate into 2 subclusters at higher resolution (inset). The “upper” and “lower” cluster were later determined to correspond to abaxial and adaxial BSC (Fig. 2), and are therefore named ^ab^BS and ^ad^BS **c.** Violin plots showing distribution of normalized mRNA counts of marker genes for cells in each cluster. Genes were canonical C4 markers (*PEP1, MDH6, CA, ME1, RBCS*) or genes that identify unique clusters (*NAAT1, TAAT*). **d.** Violin plots showing normalized mRNA levels of genes differentially expressed between ^ab^BS and BS^ad^ subclusters (*SWEET13a, b, and c*, *CC3*) and of example genes highly expressed in both clusters (*ME1*, *RBCS1*). Gene IDs are provided in Supplementary Table 4.

An estimated 7,000 protoplasts were pipetted into a 10x Chromium microfluidic chip, and single cell cDNA libraries were generated and sequenced. Gel bead-specific barcodes were used to identify mRNAs present in specific cells. After filtering the dataset to select for healthy cells, 4,035 cells with an average of 4,874 mRNA molecules per cell were analyzed. Unsupervised clustering was performed using Seurat (Butler et al., 2018) to determine the relationship between mRNA expression profiles in PCA space, ultimately represented in a two-dimensional UMAP plot (Fig. 1b). Cell identities were assigned to the clusters based on established marker genes for different cell types (Supplementary Table 1). We obtained six clusters, five of which formed a large supercluster that all had mesophyll identities, and one separate cluster corresponding to bundle sheath identity (Fig. 1c). The distribution of marker genes was consistent with the roles of mesophyll and BS cells in C4 photosynthesis (Supplementary Fig. 2, Supplementary Table 2). The ratio of mesophyll to BSC was ~75:1, indicating a low efficiency of BSC retrieval. To our surprise, no vascular cells were recovered. The BS cluster was further divided into two subclusters, the “upper” and “lower” subclusters, which later were assigned as abaxial (^ab^BS) and adaxial (^ad^BS) bundle sheath cells (Fig. 1d, Fig. 2) (see below).

**Figure 2.**
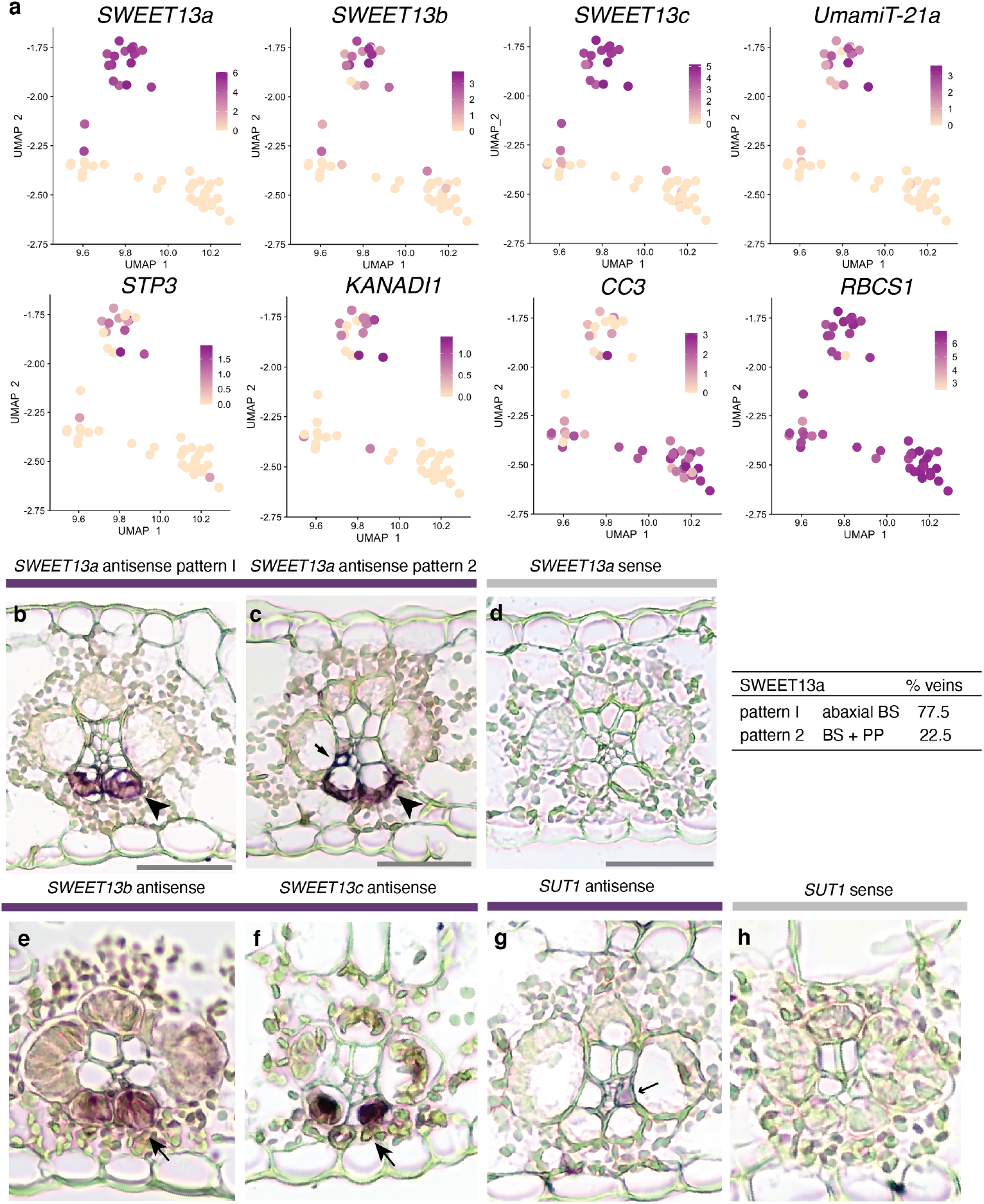
Abaxial BS cluster is enriched for genes encoding transport proteins. Feature plots show normalized levels of mRNAs for genes differentially expressed between the two clusters of bundle sheath cells plotted in UMAP space. **b - f.** *In situ* hybridization of *SWEET13a*, *SWEET13b*, and *SWEET13c.* Rank-2 intermediate veins from sections hybridized with antisense probes for *SWEET13a*, *SWEET13b*, and *SWEET13c* showed mRNA localization of three SWEET13 genes was largely limited to abaxial bundle sheath cells. **b - c.** *SWEET13a* mRNA localization was predominantly in ^ab^BS cells in the majority of veins (77.5%) and to PP in a subset of veins (22.5%) (n = 824). **d.** No staining was visible after hybridization with *SWEET13a* sense probe. **e – f.** *SWEET13b* and *SWEET13c* probes showed signal predominantly in ^ab^BS cells. **g.** *SUT1* mRNA is localized to a vascular cell which is likely a companion cell in rank-2 intermediate veins (arrow). **h.** No staining visible after hybridization with *SUT1* sense probe. See Supplementary Figure 5 for intermediate rank-1 and major veins. All error bars are 100μm.

### Mesophyll and bundle sheath clusters show canonical expression of C4 photosynthesis genes

Unsupervised clustering resulted in five clusters of mesophyll cells based on the presence of canonical C4 marker genes. At first sight the presence of five clusters was surprising. Cluster MS1 included most mesophyll cells and likely represents the core mesophyll. MS1 was enriched in photosynthetic processes. MS2 was enriched for the GO terms triose phosphate transport (GO:0035436; GO:0015717), nucleic acid metabolic process, immune system process (GO:0002376) and RNA metabolic process (GO:0016070) (Supplementary Table 1, 3). MS3 and MS4 contained high levels of rRNA transcripts. rRNA was not removed before sample preparation nor were cells with higher rRNA transcript levels, as rRNA levels can vary between cells. Whether these rRNA-enriched clusters represent biologically relevant cell populations in the leaf or are due to artifacts was not further evaluated. An additional subcluster, MS5, had an apparent mesophyll identity, but was clearly separated from the other mesophyll clusters. The main determinants for this separate clustering were iron/metal-related processes (Supplementary Text). Due to the focus on BSCs, localization and biological relevance of MS5 was not characterized further. Our clustering data are supported by the presence of transcripts for the glutamine synthetase *GLN4* (corresponding to *gln1-3* and protein GS1-3) in the MS1-4 cells, consistent with previous *in situ* and immunolocalization data that detected glutamine synthetase specifically in the mesophyll (Supplementary Fig. 3) (Martin et al., 2006).

It may be interesting to explore further whether the subclustering of mesophyll cells represents developmental trajectories or physiological differences. We did not identify an obvious pattern that could be attributed to, for example, dorsoventral patterning due to developmental gradients or due to changes in light properties as it passes through the leaf. Similar observations regarding the presence of multiple mesophyll clusters were made for scRNA-seq analyses of Arabidopsis leaves, however, it was not possible to assign palisade and spongy parenchyma to any of the mesophyll clusters (Kim et al., 2020).

### Both subsets of bundle sheath cells express canonical C4 photosynthesis genes

In maize, photosynthesis is partitioned between the mesophyll and bundle sheath cells (Li et al., 2010; Chang et al., 2012; Friso et al., 2010), allowing us to differentiate between these cell populations based on their mRNA profiles. Maize leaves utilize an NADP-ME-dependent C4 pathway, and mesophyll cells and bundle sheath cells must exchange intermediate species via plasmodesmata, with specific enzymes highly upregulated in one cell type or the other. To identify the clusters, we selected several key marker genes that are known to be differentially expressed in either mesophyll or bundle sheath.

Most of the genes involved in primary carbon metabolism, which showed significant differential expression in the 2010 proteomics survey of MS and BS chloroplasts (Friso et al., 2010), were expressed in the expected cell types in our dataset (Supplementary Fig. 2, Supplementary Table 1, 2), except for the predicted chloroplast envelope transporter *TPT* and the Calvin cycle enzyme *PGK2*, which were both expressed nearly equally in both MS and BS clusters. For example, transcripts of pyruvate orthophosphate dikinase 2 (*PDK2*) were almost exclusively found in BSCs, while *PDK1* transcript levels were high in mesophyll (Supplementary Fig. 2, Supplementary Table 1, 2), in agreement with both proteomics data (Friso et al., 2010) and their corresponding contributions to C4 photosynthesis (Chastain et al., 2017). All cells in the BSC subcluster showed high and specific expression of the canonical C4 photosynthesis-related genes that function in the BSCs, including *RBCS*, *ME1*, and *PCK1* (Supplementary Fig. 2, Supplementary Table 2). The Rubisco small chain mRNAs (*RBCS1*, *RBCS2*) were equally distributed in both abaxial and adaxial BS subclusters (Fig. 1). The bundle sheath-specific glutamine synthetase *GLN3* (corresponding to gene *gln1-4* and protein GS1-4) was found in both BS subclusters, consistent with *in situ* and immunolocalization data that showed no evidence for a specific pattern for GS1-4 in the BS (Supplementary Fig. 3) (Martin et al., 2006).

### Abaxial BS cluster is enriched for genes encoding transport proteins

The two BS cell subclusters (Fig. 1d) could either represent developmental trajectories along the leaf axis, BS cells from the three different vein classes (major vein or rank-l or rank-2 intermediate veins), different physiological states, or dorsoventral patterning. While the majority of mRNAs corresponded to BS identity, only 5 genes appeared specific to ^ad^BS (the “lower subcluster”) and 39 to ^ab^BS (the “upper” subcluster) (Table 1). Surprisingly, among the genes with the highest difference between the two BS subclusters were the three *SWEET13a*, *b*, and *c* paralogs. SWEETs are uniporters, and maize SWEET13a, b and c function as sucrose transporters.

**Table 1.**
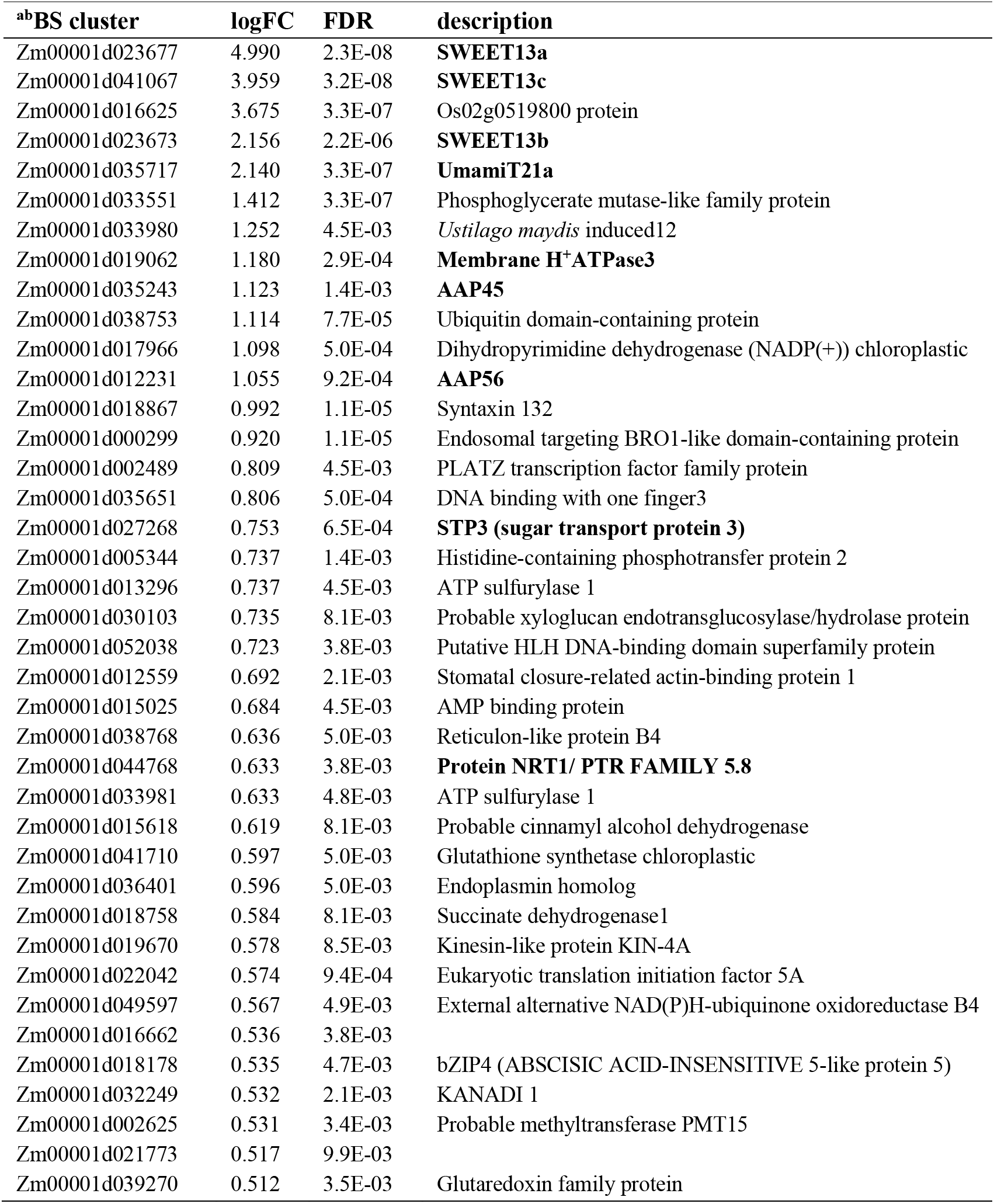
Genes of interest differentially expressed between clusters ^ab^BS and ^ad^BS. 9 out of 39 genes specific to the abBS cluster are specific to transmembrane transport (bold). Criteria for inclusion were average log fold change > 0.5 for all cells in subcluster and FDR-adjusted p-value < .01. The ^ab^BSC specificity was validated for three genes, *SWEET13a*, *b* and *c*. Whether genes with lower FDR-adjusted p-values also show high specificity will require experimental validation.

Similar to their orthologs AtSWEET11 and 12 from Arabidopsis, in maize leaves SWEET13a, b, and c are critical for phloem loading of sucrose (Bezrutczyk et al., 2018), though it remained unknown if the maize SWEET13s were co-expressed in the same cells and in which cell types they function. All three *SWEET13* paralogs were present in BSCs in maize, while the Arabidopsis orthologs *AtSWEET11*, *12* and *13* are specifically expressed in PP (Kim et al., 2020). This observation is consistent with the qRT-PCR results performed during the optimization of the protoplast protocol, which detected *SWEET13* transcripts, but no vascular cells were recovered by scRNA-seq. The “upper” BS subcluster, ^ab^BS, showed a striking enrichment for transport proteins, with 9 of 39 ^ab^BS-specific genes involved in transport (Table 1). Importantly, this included not only *SWEET13a*, *b*, and *c*, but also an STP hexose transporter (STP3), the amino acid efflux transporter *UmamiTT21a* (Supplementary Fig. 4), two members of the H^+^/amino acid symporter family, *AAP56* and *AAP45* (Deng, 2014), a member of the nitrate peptide transport family, and the H^+^-ATPase AHA3 (Table 1). Notably, in Arabidopsis, transcripts for the closest Arabidopsis homolog of Zm*UmamiT21a*, *AtUmamiT21*, were enriched in PP (Fig. 3) and co-expressed with *AtSWEET11* and *12* in Arabidopsis. On the basis of the presence of SWEETs, UmamiT21 and other ^ab^BS-enriched candidates, one may speculate that the transcription factors that in Arabidopsis are involved in PP identity have been recruited in maize to ^ab^BS to drive the unique set of genes expressed in ^ab^BS. *DOF3* (*DNA binding with one finger3*) is a transcription factor implicated in the control of SWEET gene expression in rice (Wu et al., 2018) which shows ^ab^BS-specific expression in this dataset; two other transcription factors, *bZIP4* (*ABA-insensitive 5-like protein*) and *MYB25* (just below p-value cutoff), were enriched in ^ab^BS. While ^ab^BS-enriched, *bZIP* and *MYB25* were not BS-specific, but only sparsely expressed in mesophyll cells (Supplementary Figure 3).

**Figure 3.**
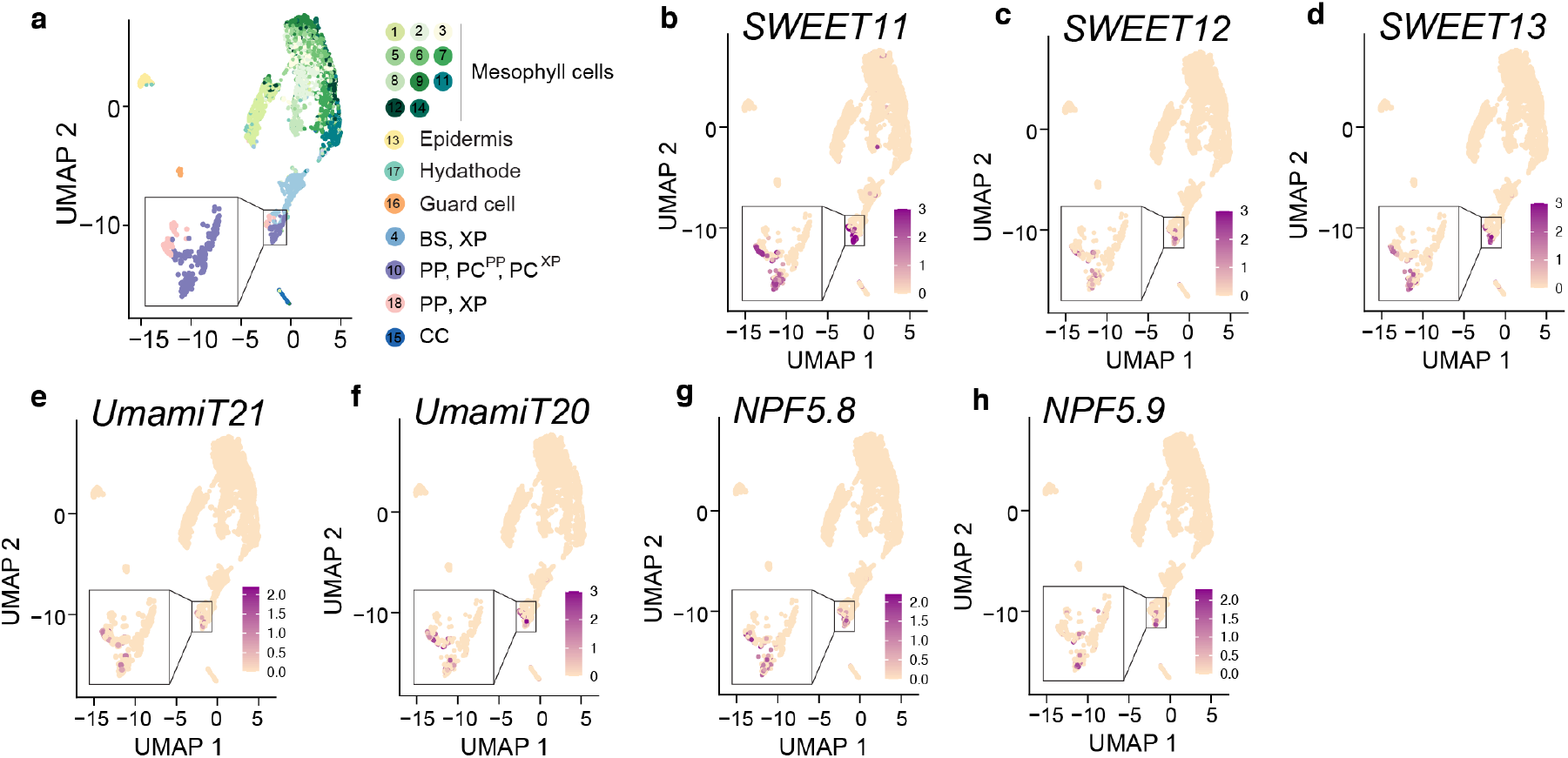
Arabidopsis homologs of maize ^ab^BS-specific genes are expressed in Arabidopsis phloem parenchyma. Maize transporters showing ^ab^BS mRNA enrichment are homologous to many Arabidopsis transporters with Arabidopsis phloem parenchyma mRNA enrichment. **a.** UMAP plot showing 2D representation of cell relationships in multidimensional space for single-cell sequencing of Arabidopsis leaf. Clusters are indicated by colors to the right of the UMAP plot. Feature plots show normalized levels of mRNA transcripts for Arabidopsis transport proteins homologous to abBS transport proteins. **b-d.** *AtSWEET11* (AT3G48740), *12* (AT5G23660), and *13* (AT5G50800) are orthologs to *ZmSWEET13a*, *b*, and *c*. **e.** *AtUmamiT21* (AT5G64700) is homologous to *ZmUmamiT21a*. **f.** *AtUmamiT20* (AT4G08290) is homologous to *ZmUmamiT21a* (Supplementary Fig. 4). g-h. *AtNPF5.8* (AT5G14940)_*and AtNPF5.9* (AT3G01350) are homologous to *ZmNRT1*.

Other BSC-specific genes such as *RBCS1* showed equal transcript distribution across all BS cells, excluding the possibility of an artefact, e.g., a gradient of cells differing in UMI counts. This included *UmamiT20a*, which was BS-enriched but equally expressed across both BS subclusters. The lack of specificity of many genes for subsets of BSC is consistent with published data from *in situ* hybridization and immunolocalization of *RBCS1* and glutamine synthetase, both of which did not show dorsoventral patterning (Langdale et al., 1988a; Martin et al., 2006; Langdale et al., 1988b).

To test the hypothesis that the two BS subclusters may represent spatially discrete BSC populations, *in situ* hybridization was used to localize the mRNAs of the *SWEET13a, b*, and *c* genes (Fig. 2b-f, Supplementary Fig. 5). Notably, transcripts of all three *SWEET13* genes were specifically detected in the two abaxial bundle sheath cells (^ab^BSCs) adjacent to the phloem. Additionally, *SWEET13a, b*, and *c* transcripts localized to the PP (Fig. 2c, Supplementary Fig. 5) similar to *AtSWEET11*, *12*, and *13* (Kim et al., 2020; Chen et al., 2012). Using three independent sets of probes, transcripts of all three SWEET13s were almost exclusively found in the ^ab^BS of rank-2 intermediate veins, which are a special adaptation of C4 monocots (Langdale et al., 1988a) and serve as the main sites of phloem loading (Fritz et al., 1989, 1983). *SWEET13a* was detected in both ^ab^BS and PP in about 23% of the rank-2 intermediate veins. Thus, in the veins that are the main loading sites, sucrose efflux towards the SE/CC for phloem loading must occur predominantly from the ^ab^BS into the apoplasm towards the phloem, and only to a smaller extent by direct release at the SE/CC from PP. Rank-1 intermediate veins seemed to have a more balanced distribution of *SWEET13a* between ^ab^BS and PP. In major veins, SWEET13 transcripts were also present in the medial vascular parenchyma, and the main path appeared to be through release from PP.

Since protein abundance does not always correlate with mRNA levels (Walley et al., 2016), we evaluated the cell specificity of the SWEET13a protein. Maize lines were generated that stably expressed translational GUS reporter fusions (*SWEET13a-GUS*) comprising 6 kb upstream of the ATG and all exons and introns. Six transgenic lines from three independent transformation events showed consistent localization of the SWEET13a-GUS fusion protein in the ^ab^BS and phloem parenchyma of rank-1 and -2 intermediate veins, and in the PP of the major veins (Fig. 4; Supplementary Fig. 5). In summary, *in situ* hybridization and immunolocalization showed that *SWEET13a, b, and c* transcripts and SWEET13a protein were not only found in the PP of maize, but also in a subset of BSCs, specifically the ^ab^BS. This is different from the cellular expression of their orthologs in the dicot Arabidopsis, suggesting that an additional sucrose phloem loading pathway evolved in C4 monocots (Langdale et al., 1988a).

**Figure 4.**
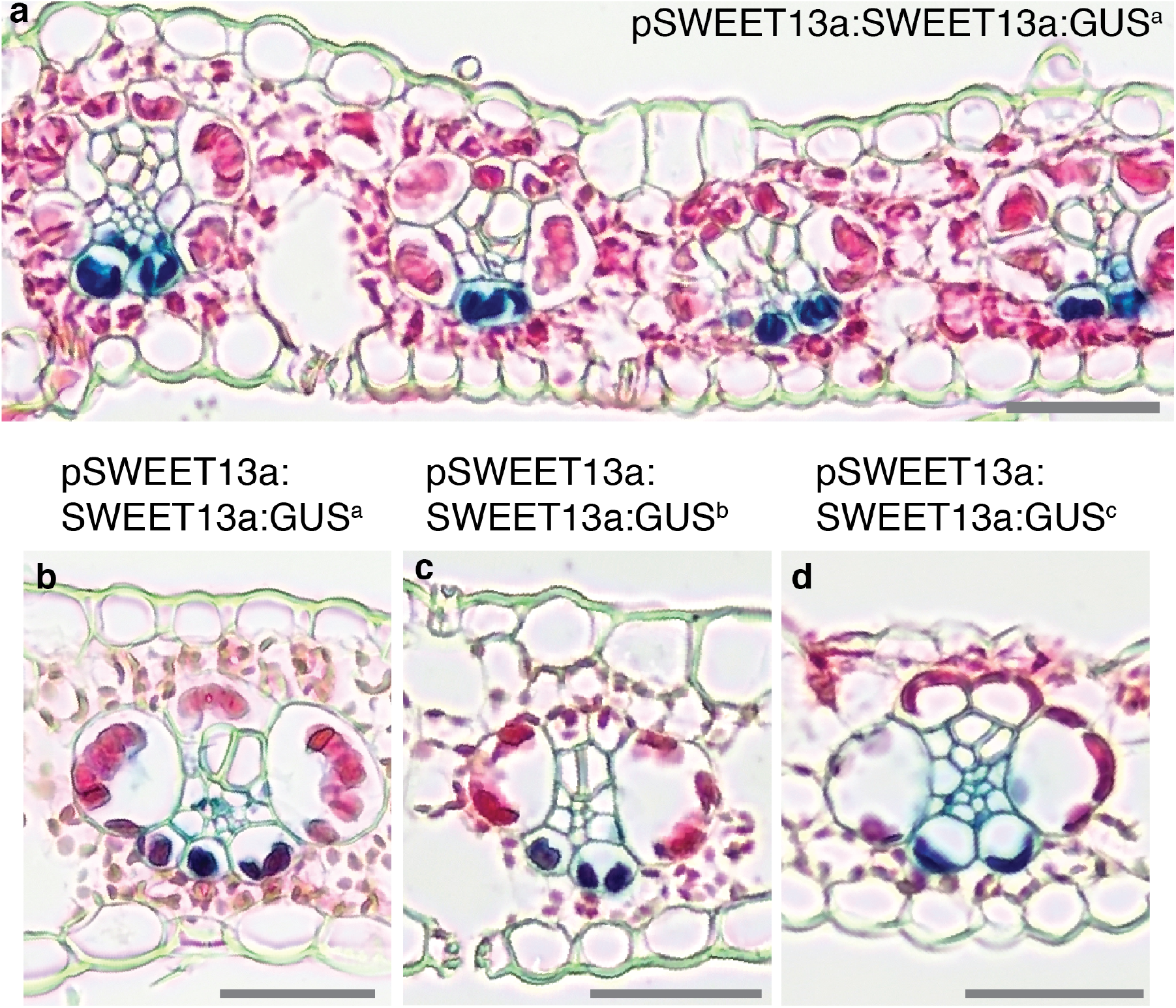
SWEET13a protein is localized to abaxial BSC of rank-2 veins. GUS histochemistry in leaves of maize lines transformed with the translational fusion construct pSWEET13a:SWEET13a-GUS. **a.** Chloro-bromoindigo precipitate (blue) is detected specifically in abaxial bundle sheath cells of maize plants transformed with pSWEET13a:SWEET13a:GUSplus. **b - d**. Three independent transformation events (a, b, and c: pSWEET13a:SWEET13a:GUS^a, b,^ and ^c^) showed similar expression patterns in rank-2 intermediate veins, rank-1 intermediate veins, and major veins (for rank-1 and major veins see Supplementary Fig. 5). **b.** Line “a,” **c.** Line “b,” **d.** Line “c.” Sections are counterstained with Eosin-Y; scale bars are 100 μm.

### *SWEET13a*- *c* and *SUT1* sucrose transporters are expressed in complementary cell types

Sucrose released from cells by SWEETs is taken up actively into the SE/CC by SUT1 H^+^/sucrose symporters (Riesmeier et al., 1994; Bürkle et al., 1998; Gottwald et al., 2000; Slewinski et al., 2009). To directly compare SWEET13 and SUT localization, *in situ* hybridization was performed in parallel using the same method from leaves at the same stages of development. *SUT1* RNA was typically found in one or two cells in the phloem, which most likely represent companion cells, where it is responsible for phloem loading. In rank-1 and major veins, *SUT1* mRNA was also detected in the medial vascular parenchyma, where it likely contributes to sucrose retrieval (Heyser et al., 1977). In our experiments, *SUT1* transcripts were not detected in bundle sheath cells, consistent with *SUT1* expression below the detection limit in of our BSC single cell dataset (Fig. 2g-h; Supplementary Fig. 5).

### Abaxial BS transcripts are co-regulated during the sink-source transition

In Arabidopsis, many PP-specific genes were found to also be co-regulated (Kim et al., 2020). We therefore tested whether several of the transporter genes identified in ^ab^BS might also be co-regulated. *SUT1* H^+^/sucrose symporter genes are typically lowly expressed in young net-importing leaves and induced during the sink to source transition (Bürkle et al., 1998; Riesmeier et al., 1993). RNA was extracted from different segments of leaf 3 of V2 plants, in which the tip had transitioned to source while the base was still in the sink state (Tausta et al., 2014) (Fig. 5a) for qRT-PCR. *SWEET13a*, and *UmamiT21, AAP45*, and *SUT1* had transcript levels that were 115-, 34-, 23-, and 10-fold higher, respectively, than in the tip of leaf 3 (source) compared to the base (sink) (Fig. 5a, b). SWEET13a protein levels were also higher in source regions of the leaf (Fig. 5c). SWEET13a was not detected in stem sections near the base of the plant, which contain whorls of developing leaves, nor in root tip (Fig. 5d-f). In leaves, SWEET13a protein was not detectable in tissue other than the tip of leaf 3, consistent with its role in phloem loading in source leaves The co-regulation of ^ab^BS-enriched genes during the developmental transition of leaves not only links them to transfer of nutrients to the phloem, but also indicates that they are all controlled by the same regulatory system.

**Figure 5.**
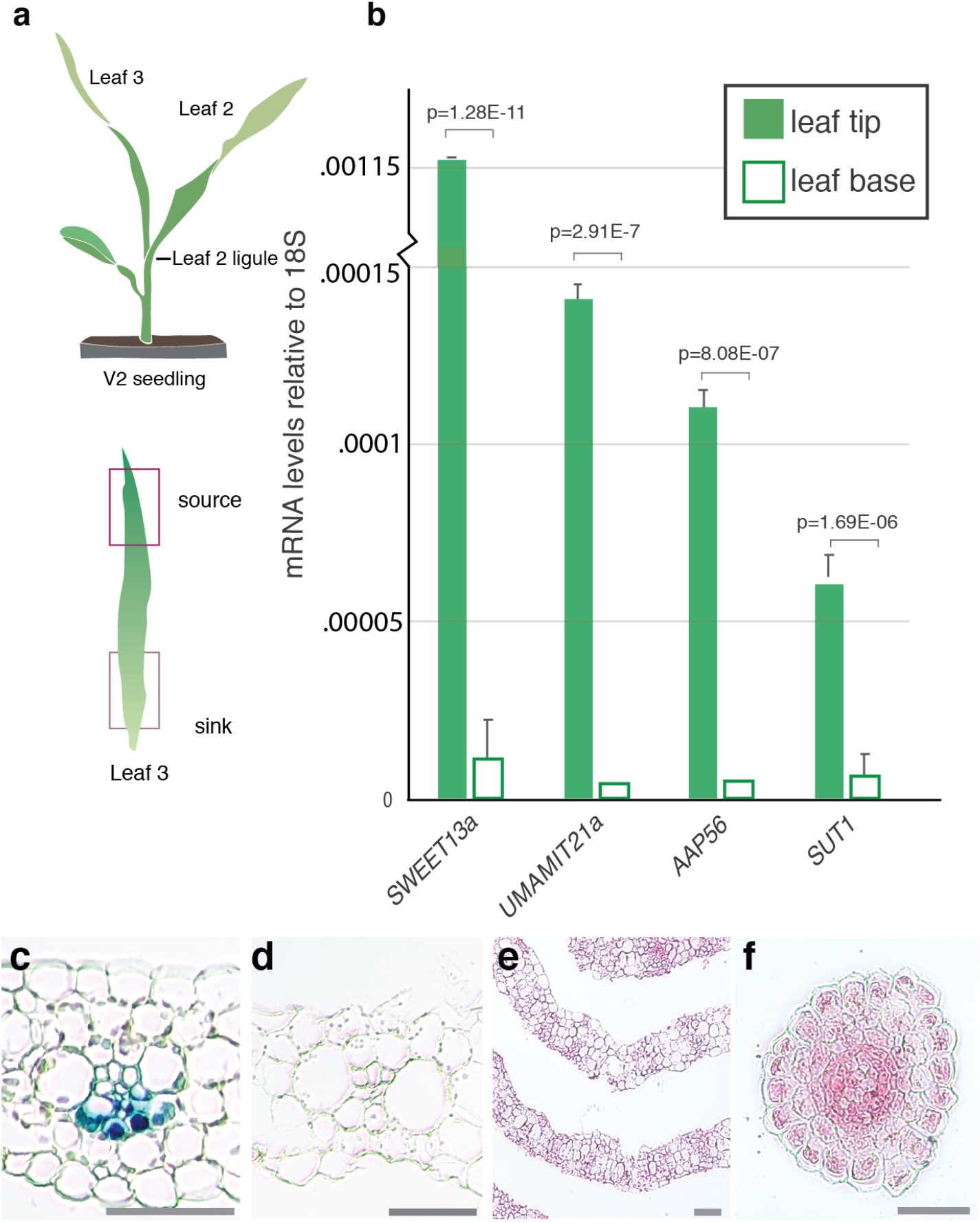
Abaxial BS transcripts are co-regulated during the sink-source transition. **a.** Tissue selected for qRT-PCR: V2-stage seedling with source and sink tissue highlighted; base of leaf 3 is still in the whorl and is net-sink tissue (Li et al., 2010). **b.** 18S-normalized mRNA levels of ^ab^BS-specific transport proteins (*SWEET13a*, *UmamiT21a*, *AAP45*) and *SUT1* in source (tip) and sink (base) tissue. Values are average of three technical (qRT-PCR) replicates of three pools of two plants; error bars represent SEM. *Students two-tailed paired t-test values shown. Independent repeats confirmed the data. **c.** pSWEET13a:SWEET13a:GUSplus transformed B73 seedling segments after 12-48 h incubation in GUS staining solution. V2 leaf 3 tip (12 h), **d.** leaf 3 sheath (48 h), **e.** stem cross section 1 cm above soil (48 h), **f.** cross section across root tip (48 h), Of these, only the tip of leaf 3 (source) showed chloro-bromoindigo precipitate indicative of SWEET13a expression. Scale bars: 100 μm.

## DISCUSSION

While single cell sequencing was successfully used to identify the transcriptomes of vascular cells in Arabidopsis (Kim et al., 2020), the suberin-lignin barrier surrounding the bundle sheath of maize leaves prevented access to vascular cell transcriptomes in maize. With an optimized protocol, BSC were released and could be identified based on a broad range of known marker genes. BSC separated into two subclusters. mRNA for BSC markers such as *RBCS* and GS1-4 were present at equal levels in both subclusters, whereas others were specifically enriched in one of the two subclusters. Because only moderately and highly expressed mRNAs are captured with droplet-based scRNA-seq protocols such as 10x Chromium, we cannot exclude that transcripts that appear to be specific are present at lower levels in the other cell types. A major surprise was the finding that mRNA for all three *SWEET13* paralogs was present in BSCs, in clear contrast to the distribution of their orthologs in Arabidopsis (Chen et al., 2012). Since barley BSCs seem clearly differentiated, with S-type cells suspected to represent a preferential site of sucrose transfer into the phloem (Williams et al., 1989), we tested whether *SWEET13a*, *b*, and *c* mRNA would be present in adaxial and lateral BSCs. To our surprise, *in situ* hybridization and the analysis of translational GUS fusions showed that all three SWEET13s were preferentially expressed in the ^ab^BS cells of rank-2 intermediate veins, which are considered the main sites of phloem loading in maize. In maize, these two abaxial BSCs are smaller compared to the medial BSCs (Bosabalidis et al., 1994). In rank-2 intermediate veins of maize, it appears that the ^ab^BS cells may have recruited sucrose-transporting *SWEETs* to export sucrose toward the abaxially localized phloem (Fig. 6). This presents a unique pathway, in which BSCs likely export photosynthetically derived sucrose to the apoplasm of the phloem on the abaxial side of the leaf. Rank-2 veins are thought to be an emergent phenomenon of C4 grasses (Sedelnikova et al., 2018). Rank-2 veins increase the ratio of BS to MS cells, the vein density, and the capacity for nutrient transport. They appear to be the main path for sucrose phloem loading. It is thus conceivable that the unique phloem loading pathway coevolved with the evolution of the rank-2 intermediate veins.

**Figure 6.**
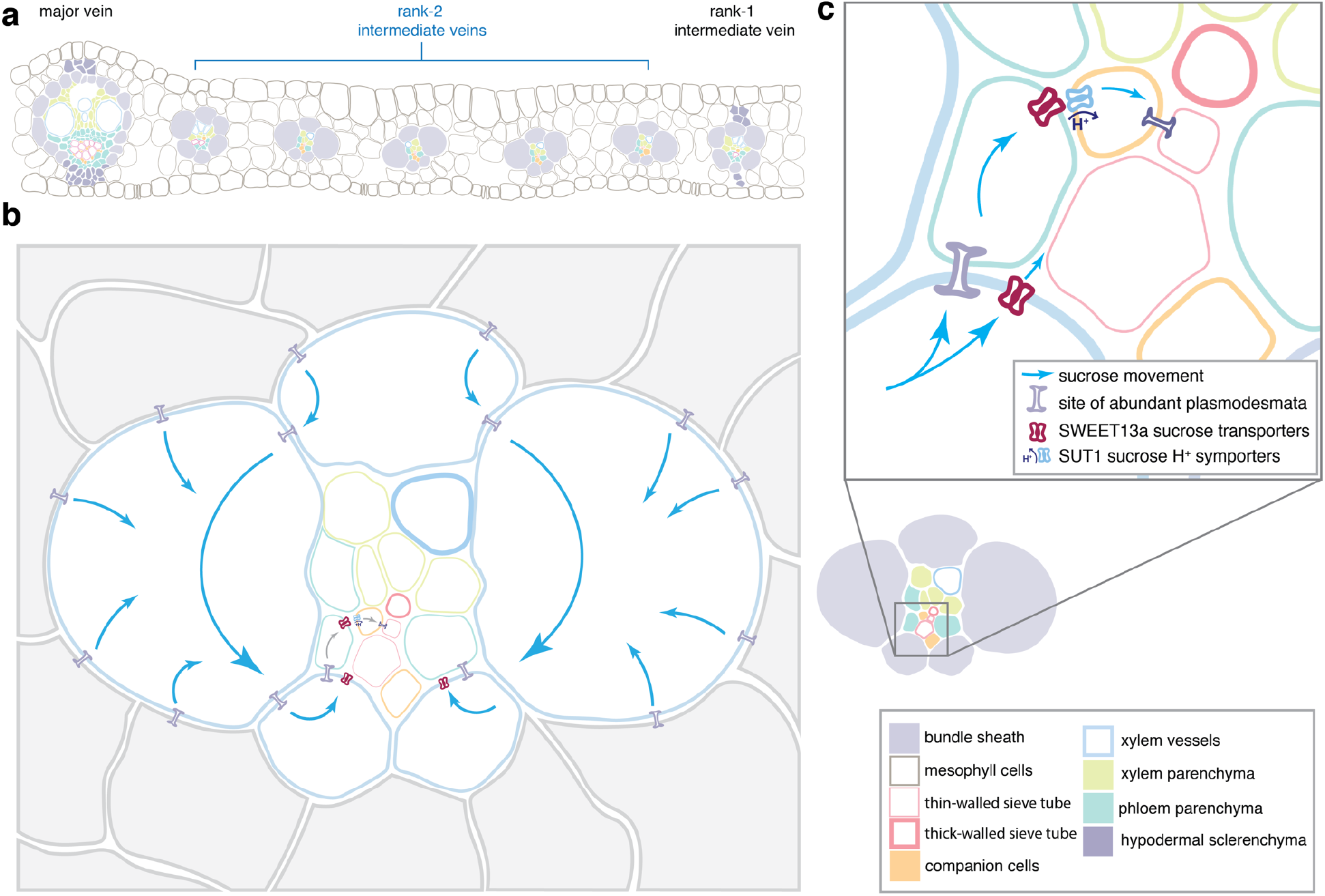
Phloem loading occurs via the ^ab^BS in maize. **a.** Example of arrangement and relative numbers of major veins, rank-1 intermediate veins, and rank-2 intermediate veins in a mature maize leaf. Note that rank-1 intermediate veins are distinguished from rank-2 by the presence of hypodermal sclerenchyma. **b.** A rank-2 intermediate vein surrounded by bundle sheath (blue outline) and mesophyll (grey) cells. Sucrose movement down its concentration gradient is indicated by blue arrows. **b.** Inset panel shows detail of sucrose movement from bundle sheath cells into apoplasm via SWEET13 transporters, or to phloem parenchyma (teal) via plasmodesmata, where it is then effluxed to the apoplasmic space by SWEETs. Sucrose in the apoplasm is taken up by SUT1 into the companion cells-sieve element (orange, pink) complex for long distance transport.

Given the findings of Williams *et al* (Williams et al., 1989), which indicate that barley uses adaxial and medial BSCs for phloem loading, our results suggest that the two species use distinct sets of BSCs for transferring sucrose from the BS to the phloem. This may be generalizable to other C3 and C4 species, and it will be interesting to explore whether *SWEETs* are also present in medial BSCs of barley. ^ab^BS and ^ad^BS transcript profiles are highly similar, possibly explaining why this differentiation of BSCs had previously not been identified.

In rank-1 intermediate and major veins, *SWEET13a, b* and *c* transcripts were also detected in cells in the vasculature that most likely correspond to phloem parenchyma, thus similar to the canonical pathway in Arabidopsis (Chen et al., 2012; Cayla et al., 2019). Phylogenetic and functional analyses had shown that the PP transporters AtSWEET11 and 12 from Arabidopsis are orthologous to the three SWEET13 isoforms (Bezrutczyk et al., 2018). Subsequent to SWEET-mediated efflux, sucrose is taken up actively by SUT1/SUC2 H^+^/sucrose symporters in both maize and Arabidopsis (Gottwald et al., 2000; Slewinski et al., 2009). This is supported by *in situ* hybridization of *SUT1* in maize leaves (Supplementary Fig. 5). PP localization of *SUT1* is consistent with previously published results (Baker et al., 2016). We could not confirm previous data that indicated that SUT1 may also be expressed substantially in BSCs (Supplementary Fig. 5). Localization of *SUT1* in vascular parenchyma (VP) is consistent with a role in sucrose retrieval on the side of the phloem that faces the xylem. Our data are compatible with the presence of two distinct sites for phloem efflux in maize leaves, one from ^ab^BS and a more standard path from PP (Fig. 6). This unique pathway may be a specific adaptation of maize leaves in the context of C4 photosynthesis, to provide higher rates of sucrose flux towards the phloem. No doubt, *SWEET13a, b* and *c* are key players for phloem loading, though at present we cannot assess the relative contribution of this new efflux step. This model could be tested by inhibiting SWEET activity specifically either in BSC or PP. However, since transcription factors driving expression of genes specifically in maize ^ab^BS or PP are not currently known, this hypothesis could be tested by generation of lines in which *SWEET13* mRNA levels have been repressed through BSC-specific RNAi.

Notably, transcripts for other transporter genes were also enriched in ^ab^BS. This includes UmamiT21a, a member of the UmamiT amino acid transporter family. One of the key findings from the analysis of PP in Arabidopsis by scRNA-seq was that multiple members of this family were specifically expressed in PP (Fig. 3, Table 1)(Kim et al., 2020). Since they appear to play roles analogous to that of SWEETs in cellular efflux of amino acids in Arabidopsis, it appears that ^ab^BS, besides having clear BSC identity, has acquired components or subnetworks of the PP identity. The co-regulation of at least some of the ^ab^BS-enriched genes further strengthens this hypothesis. Interestingly, we also observed a weak enrichment of the abaxial KANADI transcription factor (Fig. 2a). KANADI plays a key role in determining abaxial identity in leaves (Candela et al., 2008). We therefore hypothesize that one or several transcriptional regulators that are involved in the regulation of the efflux of sucrose and amino acids from PP have been brought under control of both a polarity cue and the BS identity cues in order to increase nutrient flux towards the maize phloem. It will be fascinating to identify the transcription factors that are involved in controlling the PP and BSC identities and to dissect the SWEET promoters to determine which *cis*-elements are involved in the acquisition of the ^ab^BS cell fate. Several transcriptional regulators have been identified as candidates for the induction of ^ab^BS cell fate; however, the limitations of sequencing depth in 10x Genomics Chromium droplet-based scRNA-seq preclude a comprehensive profiling of all transcriptional regulators. Methods that provide higher sensitivity may help to address this aspect. Importantly, the ^ab^BS genes identified provide unique insight into the specialized nature of this cell type.

Comparison of this phenomenon in other grasses, those that use both C3 and C4 photosynthesis, as well as a careful analysis of the evolution of rank-2 intermediate veins may provide insights into how widely distributed this mechanism is and may provide hints regarding the evolution of this regulatory rearrangement. Finally, new methods will be required to gain access to the vascular cells of maize, which are not accessible through the current methods. In summary, scRNA-seq enabled the identification of cells with a unique combination of properties on the adaxial side of the bundle sheath, cells that play key roles in C4 photosynthesis. The identification of this new property may be relevant to bioengineering of staple crops, for example, C4 rice.

## Materials Availability

Plasmids generated in this study have been deposited to Addgene under the code Plasmid #159535.

## Data and Code Availability

The raw data that support the findings of this study are available from the corresponding author upon reasonable request. All sequencing data will be deposited in the Gene Expression Omnibus GEO (www.ncbi.nlm.nih.gov/geo/) and the accession number will be updated here.

## METHOD DETAILS

### Plant growth

*Zea mays* L. B73 seeds were germinated on filter paper with deionized, distilled water in darkness at 22-25 °C and transferred to soil upon coleoptile emergence. Plants were subsequently grown at 28-30 °C in a greenhouse supplemented by sodium lamps (400 μmol m^−2^ s^−1^) from 8:00-20:00. Protoplasts for scRNA-seq were generated from the last 6 cm of the distal portion of V2 leaf 2. For each pool of protoplasts tested, leaf segments from six concurrently grown plants were used. *In situ* hybridization was performed on sections taken from the distal portion of leaf 2 of V2 plants, distal portion of leaf 5 of V4 plants, and distal portion of the flag leaf from VT plants, with similar results. All images shown are from V4 leaf 5. For GUS staining, tissue segments were taken 10 cm from the tip of the third leaf below the flag leaf of T0 plants at growth stage VT (mature leaf tip), 4 cm from the tip of leaf 3 of T1 V2 plants (seedling leaf tip), 12 cm from the tip of T1 V2 plants (seedling leaf base), a stem section 1 cm above soil surface of T1 V2 plant (seedling stem), or the seminal root tip from T1 V2 plant (seedling root).

### Genes analyzed here

Gene IDs are provided as Supplementary Table 4.

### Probe preparation for *in situ* hybridization

RNA was extracted from leaves of V2 B73 seedlings by phenol-chloroform extraction as previously described (Bezrutczyk et al., 2018). cDNA synthesis was performed using QuantiTect Reverse Transcription Kit (Qiagen, Hilden, Germany). cDNA was amplified for each gene (Supplementary Table 5) using Takara PrimeSTAR GXL polymerase then subcloned into pJET1.2 using CloneJet PCR cloning kit (ThermoFisher, Meerbusch, Germany). For *SWEET13a, b*, and *c*, three unique regions in the 5’- and 3’-UTRs and in the coding region with lengths of ~100 bp were selected as probe templates. The three probes specific for one of the genes were combined for detection of the respective target gene. For *SUT1*, two regions in the 5’- and 3’-UTRs, unique to *SUT1* but common to all six isoforms, were selected as probe templates. All cDNA sequence alignments were performed using Geneious R11 (https://www.geneious.com) (Supplementary Fig. 6). Probe template regions were amplified with SP6 sequences flanking the forward primers for the sense probe, and reverse primers for antisense probes (Supplementary Table 5). The MEGAscript SP6 Transcription kit (ThermoFisher) was used with a 1:2 ratio of DIG-labelled UTP:UTP to generate DIG-labelled probes. Probes were precipitated after DNAse reaction by addition of 2 mg/mL glycogen, 0.1 volume 10% acetic acid, 0.1 volume NaOAc, and 2.5 volumes ethanol and centrifuged at 4 °C at 20,000 x *g* for 30 min. Pellets were washed with 70% ethanol in DEPC-treated water, allowed to dry, and resuspended in 25 μL RNAse-free 10 mM Tris-EDTA pH 8 and 25 μL formamide.

### *In situ* hybridization

*In situ* hybridization was adapted from Jackson and Simon lab protocols (Jackson, 1992; Stahl and Simon, 2010). Leaf tip sections 1 cm in length were dissected from V2 or V5 plants into 4% paraformaldehyde, vacuum-infiltrated for 10 min and fixed overnight at 4 °C. Dehydration by ethanol series and paraplast embedding were performed as described (Malcomber and Kellogg, 2004). Sections (10 μm) were cut with a Leica RM 2155 microtome and mounted on ProbeOn Plus slides (Fisher). After deparaffinization with Histoclear and rehydration by a decreasing ethanol series, tissue was permeabilized in a 2 μg/mL proteinase K solution, washed with 0.2% glycine and 1x PBS (1.3 M NaCl, 0.07 M Na_2_HPO_4_, 0.03 M NaH_2_PO_4_) for 2 min, and re-fixed with 4% paraformaldehyde for 10 min. Slides were washed with PBS and acetylated with 0.1 M triethanolamine and acetic anhydride for 10 min, then washed and dehydrated with an increasing ethanol series. Probes for each construct were mixed (e.g. all three antisense probes for *SWEET13a*), diluted 1:50 with formamide, denatured at 95 °C for 3 min, and further diluted 1:4 with hybridization buffer (300 mM NaCl, 10mM NaH_2_PO_4_, 10mM Na_2_HPO_4_, 10mM Tris-Cl pH 6.8, 5 mM EDTA, 50% formamide, 12.5% dextran sulfate, 1.25mg/mL tRNA). Probe incubation in slide pairs was performed at 55 °C overnight. Slides were rinsed three times with 0.2x SSC pH 7 (600 mM NaCl, 60 mM sodium citrate) at 55 °C for one hour, and washed with block reagent solution (Roche), washed with BSA blocking solution (10 mg/mL BSA, 0.1 M Tris-Cl, 150 mL NaCl, 0.3% Triton X-100) for 45 min, and incubated with anti-DIG antibody (Roche) for 2 hours at 22 °C. Slides were rinsed four times with BSA block solution for 15 min each, in Buffer C (100 mM Tris pH 9.5, 50 mM MgCl_2_, 100 mM NaCl) for 15 minutes, and incubated with 50 μL NBT and 37.5 μL BCIP in 5 mL buffer C for 24-48 hours. Slides were washed with water, dehydrated with an increasing ethanol series, and mounted with Eukitt Quick-hardening mounting medium. Images were taken with an Olympus CKX53 cell culture microscope with EP50 camera. *In situ* hybridization experiments for each gene (probe combination) were performed as (at minimum) three independent experiments.

### Generation of ZmSWEET13-GUS constructs

*ZmSWEET13a* (Supplementary Table 4), including 5751 bp upstream of the start codon and 684 bp downstream of the stop codon, was isolated from B73 gDNA (Supplementary Table 5) and inserted into pJET using the CloneJET PCR cloning kit. The final construct consists of GUSplus inserted directly upstream of the *ZmSWEET13a* stop codon, preceded by a 9-alanine linker, in the Golden Gate vector pGGBb-AG, the *in silico* cloning of which was performed using Geneious R11 (Supplementary Fig. 6). The assembly of all fragments with the vector pGGBb-AG was performed using the Takara InFusion HD cloning kit, and validated by Sanger sequencing.

### Maize transformation

*Agrobacterium tumefaciens* strain EHA105 was transformed with the ZmSWEET13a:GUS vector at the Crop Genome Engineering Facility at VIB Ghent (https://www.psb.ugent.be/cgef). Transformed EHA105 carrying the respective plasmids was used to transform the inbred maize line B104 via Agrobacterium-mediated transformation of 600 immature embryos according to previously described methods (Coussens et al., 2012). In brief, callus formation was induced using auxins and transgenic cells selected over several weeks using phosphinotricin (PPT) selection. Plantlets were then regenerated on hormone-free medium, and presence of the transgene confirmed using TraitCheck (Romer Labs; Butzbach, Germany) and PCR analysis. Three independent transformation events were derived from different starting immature embryos, yielding six plants in total: three plants from event A (pSWEET13a:SWEET13a:GUSplus^a^), two from event B (pSWEET13a:SWEET13a:GUSplus^b^), and one from event C (pSWEET13a:SWEET13a:GUSplus^c^).

### GUS histochemistry

Tissue segments were taken 10 cm from the tip of the third leaf below the flag leaf of T0 plants at growth stage VT (mature leaf tip), 4 cm from the tip of leaf 3 of T1 V2 plants (seedling leaf tip), 12 cm from the tip of T1 V2 plants (seedling leaf base), on the stem 1 cm above the soil surface of T1 V2 plant (seedling stem), or from the seminal root tip of T1 V2 plant (seedling root) at 13:00 o’clock. Tissue segments were dissected into cold acetone and vacuum-infiltrated for two min, vacuum infiltrated with GUS wash buffer (20 mM EDTA, 40 mM C_6_N_6_FeK_3_, 40 mM C_6_FeK_4_N_6_, 20% methanol, 57.6 mM Na_2_HPO_4_, 42 mM NaH_2_PO_4_, 0.1% Triton X-100), and incubated with GUS wash buffer including 0.2% X-Gluc at 37°C for 1-48 hours. Sections were dehydrated in 20%, 30%, 50% ethanol for 30 min, fixed in FAA (50% ethanol, 3.7% formaldehyde, 5% acetic acid) for 30 min, and further dehydrated in 75% and 100% ethanol. Embedding was performed by incubating sections at 60 °C in tert-butyl ethanol:paraplast dilutions at 3:1, 1:1, and 1:3 ratios. Melted paraplast (100%) was changed twice daily for three days. Paraplast-embedded tissue was poured into blocks, and 10-μm sections were cut with a Leica RM 2155 microtome. Sections were mounted on SUPERFROST PLUS Gold Slides (Thermo Scientific), deparaffinized with Histoclear, and mounted with Eukitt Quick-hardening mounting medium.

### Single cell sequencing: protoplast preparation

Tissue was sampled from the distal portion of leaf 2 (from 1 cm to 7 cm, as measured from the tip) from V2 plants. This region was selected because it is thought to be non-expanding, non-differentiating source tissue based on results from the RNAseq-defined developmental transcriptome of the maize leaf (Li et al., 2010). Leaf segments were harvested at 9:00 am, and tape was applied to adaxial epidermis to stabilize the tissue, which was scored every 5 mm from the midvein to leaf edge with a razor manifold consisting of scalpel blades taped together to ensure minimum distance between scores. Tape sections were placed abaxial-side down in pretreatment solution (2 mM L-cysteine, 164 mM sorbitol) and vacuum-infiltrated for 10 min with 2 min of active pumping. Tape sections were incubated with gentle agitation (30 rpm, IKA Rocker 3D orbital shaker) for an additional 20 min in pre-treatment solution, then transferred to enzyme solution (cellulase “Onozuka” RS 1.25%, cellulase “Onozuka” R-10 1.25%, pectolyase Y-23 0.4%, macerozyme R-10 0.42%, sorbitol 0.4 M, MES 20 mM, KCL 20 mM, CaCl_2_ 10 mM, BSA 0.1%, ß-mercaptoethanol 0.06%) for 3.5 h with gentle agitation (30 rpm on orbital shaker). Protoplasts were filtered through a Corning 70-μm nylon mesh strainer and centrifuged in a round bottom tube for 1.5 min at 100 x *g*. Enzyme solution was gently removed and replaced with cold wash solution (sorbitol 0.4M, MES 20mM, KCl 20mM, CaCl_2_ 10mM, BSA 0.1%). Protoplasts were carefully resuspended in wash solution and centrifuged for 30 seconds at 100 x *g*, then strained through a 70-μm Scienceware Flowmi Cell Strainer to remove large debris. Washing solution steps were repeated four additional times to remove chloroplasts and small debris. Cell viability and concentration were quantified under an Olympus CKX53 cell culture microscope: 1 μL 0.4% Trypan blue was added to 9 μL of resuspended protoplasts in wash solution and pipetted into the chamber of a C-Chip Neubauer Improved Disposable Haemocytometer (NanoEntek; Seoul, South Korea); healthy (unstained) cells were counted. Protoplasts were resuspended to a concentration of 1,200 cells /μL. A variety of approaches to degrade the suberin-lignin-containing bundle sheath cell walls with the addition of other enzymes failed to produce healthy cells: Laccase (Sigma) and manganese peroxidase (Sigma), as well as enzymes provided by Novozymes (Copenhagen, Denmark), namely a cutinase, a fungal carbohydrase blend produced in *Aspergillus aculeatus*, a fungal beta-glucanase blend produced in *Humicola insolens*, a pectinase preparation produced in *Aspergillus*, a xylanase blend, and a multi-enzyme complex containing carbohydrases, including arabanase, cellulase, beta-glucanase, hemicellulose and xylanase, were each added to the existing protocol to a final concentration of 1-2% active enzyme weight/vol. Visual inspection of protoplasts during isolation revealed that addition of these enzymes caused protoplasts to rupture.

### Protoplast protocol optimization

468 Several variations of the above protocol were tested prior to the final protoplast preparation, and the presence of diverse cell types was verified by qRT-PCR using primers specific to *MDH6*, *ME1*, *SWEET13a*, *SWEET13b*, and *SWEET13c* (Supplementary Table 5). Briefly, RNA was extracted from protoplasts using the RNEasy Mini Kit, and first strand cDNA was synthesized using Quantitect reverse transcription kit (Qiagen, Hilden, Germany). Quantitative reverse transcription PCR (qRT-PCR) was performed using Roche LightCycler 480 SYBR Green I Master polymerase on a Stratagene Mx3000P, and relative expression of transporter genes was calculated relative to 18S and Actin using the 2^−ΔCT^ method. Modifications to the standard protoplast isolation protocol of 2 h in enzyme solution (see previous section) (Protocol 1) included doubling the concentration of enzymes in solution (Protocol 2), isolating BS strands released after 2 h followed by continued incubation of filtered BS strands in fresh enzyme solution to deplete mesophyll cells (Protocol 3), and incubating the leaf tissue in pretreatment solution (Protocol 4) (2 mM L-cysteine and164 mM sorbitol). The protocol which yielded the highest ratio of BS:MS marker genes (*ME1*: *MDH6*) included a pretreatment incubation step and 1x enzyme solution (Supplementary Fig. 1).

### Cell partitioning, library prep, and sequencing

To aim for partitioning of 7,000 cells, with the expectation that 3,500 cells would be sequenced, 6 μL of the protoplast suspension with an estimated 1,200 cells/μL was applied to the 10x Genomics Chromium microfluidic chip (Chemistry V3.0). Thereafter the standard manufacturer’s protocol was followed. Twelve cycles were used for cDNA amplification, and the completed cDNA library was quantified using an Agilent AATI Fragment Analyzer. Sequencing was performed at Novogene (Sacramento, CA, USA) on a single lane with the Hi-Seq platform and the standard PE150 sequencing parameters.

### Generation of single-cell expression matrices

Cellranger count (10x Genomics) was used to process fastq files provided by Novogene, with read 1 trimmed to 26 bp (r1-length=26), as the first 26 bp of a 10x library R1 comprise the cell barcode and UMI index, and the remaining comprises poly-A tail with no further information. A formatted reference genome was generated using Cellranger mkref using the maize B73 RefGen 4 (Jiao et al., 2017) whole genome sequence and annotation (fasta and gff3 downloaded from Ensembl B73 RefGen V4), to which reads were aligned using STAR (Dobin et al., 2013). For analysis of single cell sequencing data, see “Quantification and Statistical Analysis” section below.

### Phylogeny of UmamiT transporters

BLAST results from the seed sequence AtUmamiT12 (At2g37460) to maize (AGP v4 (Jiao et al., 2017) and barley (IBSC v2 (Mascher et al., 2017)) were combined with BioMart (Smedley et al., 2009) results and filtered for the WAT1-related protein domain (panther ID PTHR31218). Genes passing this filter were selected as UmamiT family candidate genes. Two trees were generated: one using an alignment of all known splice variants, and one with only the representative transcript, with similar results. Alignment was performed in MEGA7 (Kumar et al., 2016) using MUSCLE with the following parameters: gap open penalty −2.9, gap extend penalty 0, hydrophobicity multiplier 1.2, max iterations 8, clustering method UPGMA for iteration 1, 2; UPGMB for all subsequent iterations, and lambda 24. The maximum likelihood tree was created from these alignments using IQTREE webserver (Trifinopoulos et al., 2016) using the BLOSUM62 substitution model and 1000 bootstraps.

## QUANTIFICATION AND STATISTICAL ANALYSIS

### Sample selection for scRNA-Seq, qRT-PCR, and RNAseq

Plants chosen for protoplast release, qPCR, and RNAseq were randomly selected from among a larger number of individuals which had been grown concurrently and were at the same growth stage. True biological replicates (i.e., independently grown plants) were used as replicates for statistical analyses. The number of plants per sample and number of replicates is given in the Figure legends or in specific methods sections. To ensure reproducibility, the plants used in successive experiments were grown in the same greenhouse under controlled conditions. Samples for a given experiment were taken at the same developmental stage, at the same time of day.

### Dimensionality reduction and cell clustering

The Seurat R package (v3.1)(Butler et al., 2018) was used for dimensionality reduction analysis and dataset filtering. To remove cells with low mRNA count (nFeature_RNA) and doublets, as well as damaged cells with high chloroplast (pt) or mitochondria (mt) genome-derived transcripts, cells were filtered (percent.pt <4 & percent.mt <0.75 & nFeature_RNA >1800 & nFeature_RNA <7000). Normalization, scaling, and variable feature detection were performed using SCTransform (Hafemeister and Satija, 2019). Cells were clustered using FindNeighbors to create a K-nearest neighbors graph using the first 50 principle components. FindClusters was used to iteratively group cells using a resolution of 0.2 or 23. These clusters were used as input for non-linear dimensional reduction using Uniform Manifold Approximation and Projection (UMAP)(McInnes et al., 2018).

### Differential gene expression analysis across clusters

Genes differentially expressed across clusters or subclusters were identified by comparing average normalized mRNA counts in cells of a given cluster to that of cells in all other clusters using the Seurat function FindMarkers. Genes with an FDR corrected P-value <0.05 and an average logFC >0.5 were considered marker genes.

### Identification of cluster identities

Canonical C4 photosynthesis-related genes were used as markers to define MSC and BSC clusters (Denton et al., 2017) (Supplementary Table 1). The cluster identified as BSC was subdivided into two subclusters when FindClusters was applied with a resolution of 23, and differential gene expression analysis was performed on these two subclusters with FindMarkers (for subclusters: logFC >0.5, FDR ≤0.01).

### qRT-PCR of transporter genes in seedling leaf

Leaf segments were harvested from the distal and proximal end (tip and base) of leaf 3 of early V2 plants at 13:00. Tissue was ground in liquid nitrogen and RNA extracted as previously described (Bezrutczyk et al., 2018). First strand cDNA was synthesized using Quantitect reverse transcription kit (Qiagen). qRT-PCR to determine relative mRNA levels was performed using a Stratagene Mx3000P with primers for *18S, Actin, SWEET13a, 13b, 13c, SUT1, UmamiT21a*, and *STP3* (Supplementary Table 5). Relative expression of transporter genes was calculated relative to *18S* and *Actin* using the 2^−ΔCT^ method for quantification, with similar results. Values shown in Figure 5b are the average of three technical (qRT-PCR) replicates of three pools of two plants; error bars represent SEM. Students two-tailed paired t-test values are shown. Two independent repeats confirmed the data.

### Gene Ontology term analysis for mesophyll clusters

Marker genes for each of the five mesophyll clusters (LogFC >0.5; FDR-adjusted p-value <0.05) were used as input for Gene Ontology (Ashburner et al., 2000; The Gene Ontology Consortium, 2018) analysis via the online portal GO Gene Ontology database (doi: 10.5281/zenodo.3727280; released 2020-03-23). GO terms with FDR-corrected p-values <0.05 can be found in Supplementary Table 3.

### Protoplast and bulk leaf tissue RNAseq

Protoplasts were generated according to the previously described method. Whole leaf tissue from sibling plants was ground in liquid nitrogen at the time leaf tissue was harvested for protoplast isolation. RNA from two pools of protoplasts made from four leaves each (P1 and P2), and two pools of four whole leaf segments each (L1 and L2) was extracted using the RNEasy Mini Kit (Qiagen), and four cDNA libraries were generated using the NEBNext Ultra DNA Library Prep Kit for Illumina (New England Biolabs, MA, USA) with modifications to select for 250-500 bp fragments. Sequencing of the four libraries was performed at Novogene (Sacramento, CA, USA) on a single lane with the Hi-Seq platform and the standard PE150 sequencing parameters. Reads were analyzed using a custom implementation of the Wyseq RNAseq analysis pipeline (https://github.com/astauff/WySeq). Briefly, reads were trimmed using TrimGalore (v 0.6.5) and aligned to the AGPv4 B73 reference genome using STAR (v 2.5.1b). Counts were generated using Subread featureCounts (v 2.0.1), and differential expression was analyzed using the R-packages EdgeR (v3.30.3) and limma (v 3.44.3) using trimmed mean M-value (TMM) normalization factors. Reads corresponding to BSC-specific genes and MS-specific genes were normalized separately to compensate for the expected difference in cell populations represented in protoplasts and whole leaf tissue. Genes were filtered to remove those with a coefficient of variation >75^th^ percentile within replicate groups prior to correlation analysis. Pearson correlation (Supplementary Fig. 7) and differentially expressed genes specific to BS and MS-cells (logFC > 1 or < −1) are presented (Supplementary Table 6). None of the genes in the ^ab^BS subclusters were induced by protoplast isolation. Rather, several showed reduced mRNA levels in the protoplast sample.

## ACKNOWLEDGMENTS

We would like to thank Reinout Laureyns (VIB-UGhent) for advice regarding in situ hybridization, Kajetan Linkert (HHU) regarding GUS histochemistry, Max Blank and Tatjana Buchmann (WF lab) for cDNA library preparation, and Thomas Kleist (HHU) for advice on phylogenetic analyses. This research was supported by Deutsche Forschungsgemeinschaft (DFG, German Research Foundation) under Germany’s Excellence Strategy – EXC-2048/1 – project ID 390686111 and SFB 1208 – Project-ID 267205415, as well as the Alexander von Humboldt Professorship to WBF.

## AUTHOR CONTRIBUTIONS

Conceptualization, M.B., J.K., W.B.F.; Methodology, M.B., C.P.S.K, T.H., T.L.; Investigation, M.B, N.Z, T.L, C.P.S.K.; Resources, K.K, W.B.F.; Writing, M.B., W.B.F.; Supervision, J.K., W.B.F.

## SUPPLEMENTARY DATA

### Supplementary text

#### A subset of mesophyll cells appears to be specialized in iron metabolism

Subcluster MS5 (visible in the lower left corner of Figure 1b) shared mesophyll identity but appeared be specialized in metal accumulation and transport, as indicated by high and specific expression of four nicotianamine synthase genes, *NAS1, 2*, 9, and *10*, which are involved in iron chelation; one iron phytosiderophore transporter. *YS1* (*yellow stripe 1*); and various additional genes involved in metal transport and metabolism (Supplementary Table 1). M5 cells could either represent a cell type with a specific localization in the leaf or correspond to cells that contain different levels of iron.

**Supplementary Figure 1.**
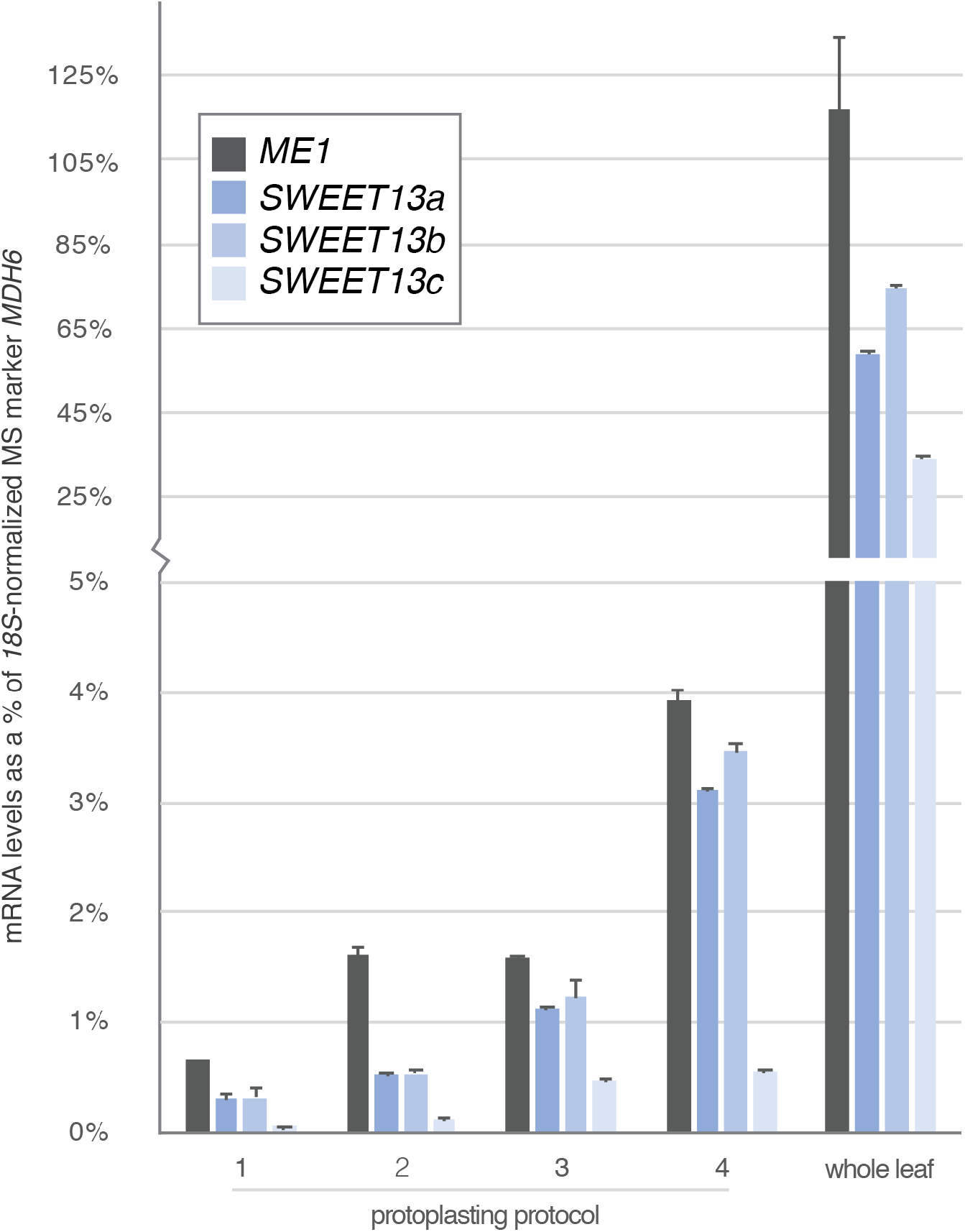
qRT-PCR of putative BSC and vascular-expressed genes as an indication of protoplast cell type diversity prior to sequencing. Normalized mRNA levels of *ME1*, *SWEET13a*, *SWEET13b*, and *SWEET13c* shown as a percentage of *18S*-normalized expression of a mesophyll marker gene, *NADP-malate dehydrogenase6* (*MDH6*), after different protoplasting treatments. Error bars represent SEM of technical duplicates. Protocol 1, standard enzyme cocktail (see Materials and Methods) with 3.5 h incubation. Protocol 2, doubled enzyme concentration. Protocol 3, isolated BS strands released after 2 h and continued protoplasting of filtered BS strands in fresh enzyme solution to deplete mesophyll cells. Protocol 4, incubated leaf tissue in pretreatment solution (2 mM L-cysteine and 164 mM sorbitol), which yielded the highest ratio of BS:MS marker genes.

**Supplementary Figure 2.**
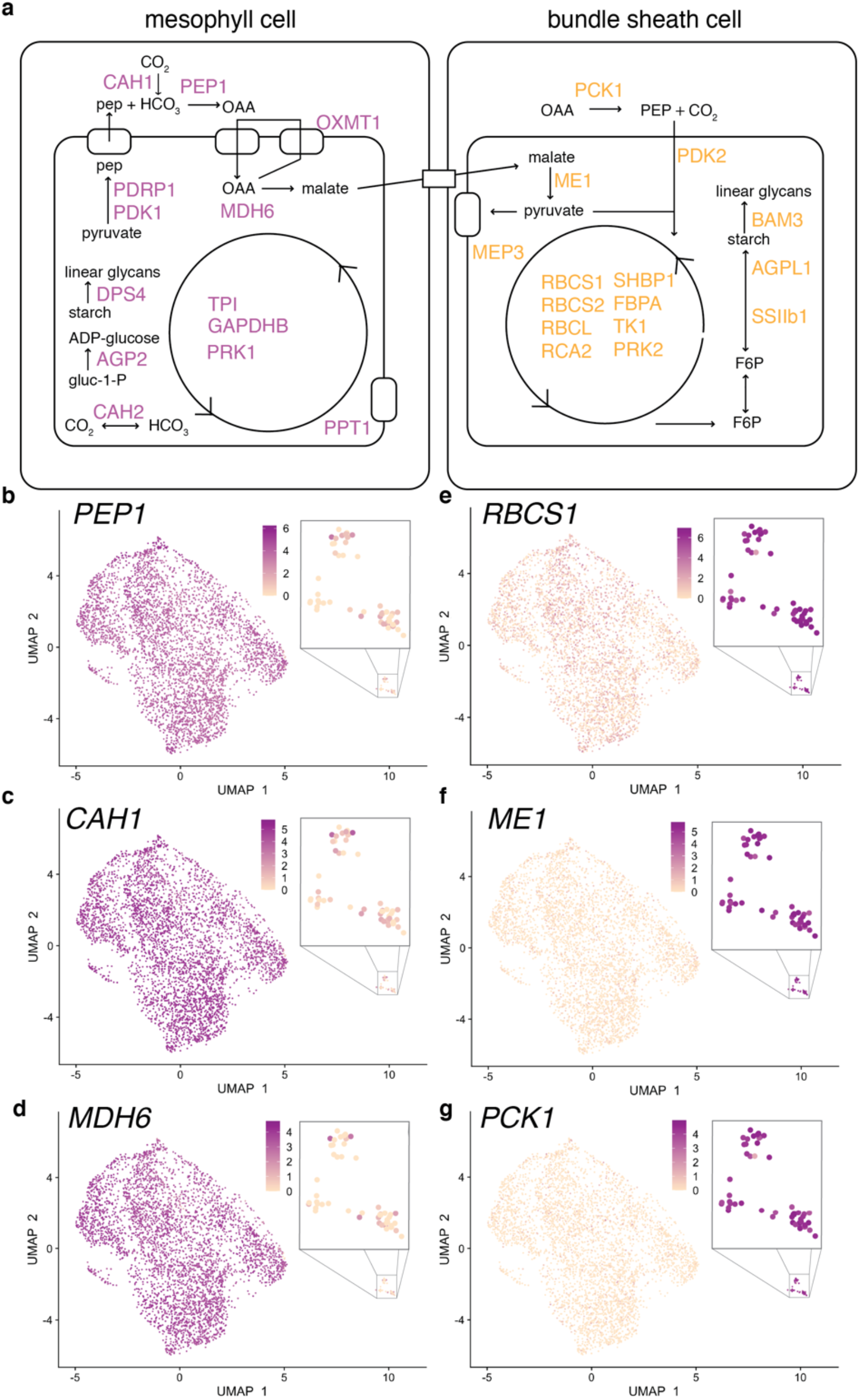
Schematic of C4 photosynthesis related genes and relative expression in BS and MS clusters. **a.** Partitioning of proteins involved in C4 photosynthesis between mesophyll and bundle sheath cells in maize(Friso et al., 2010). For all proteins displayed, mRNA expression in this scRNA-seq dataset was specific to either mesophyll or bundle sheath cells (logFC > 1.0; FDR-adjusted p-value < .05). **b**. Gene IDs, symbols, and full names are shown along with Log FC values in Supplementary Table 2. **b-g.** Feature plots show normalized levels of mRNAs for canonical C4 photosynthesis genes expressed differentially in mesophyll and bundle sheath. **b.** *PEP1* (Phosphoenolpyruvate carboxylase 1; Zm00001d046170) **c.** *CAH1* (Carbonic anhydrase 1; Zm00001d044099) **d.** *MDH6* (NADP-dependent malate dehydrogenase 6; Zm00001d031899) **e.** *RBCS1* (Ribulose bisphosphate carboxylase small subunit 1; Zm00001d052595) **f.** *ME1* (NADP-dependent malic enzyme 1; Zm00001d000316) **g.** *PCK1* (Phosphoenolpyruvate carboxykinase 1; Zm00001d028471)

**Supplementary Figure 3.**
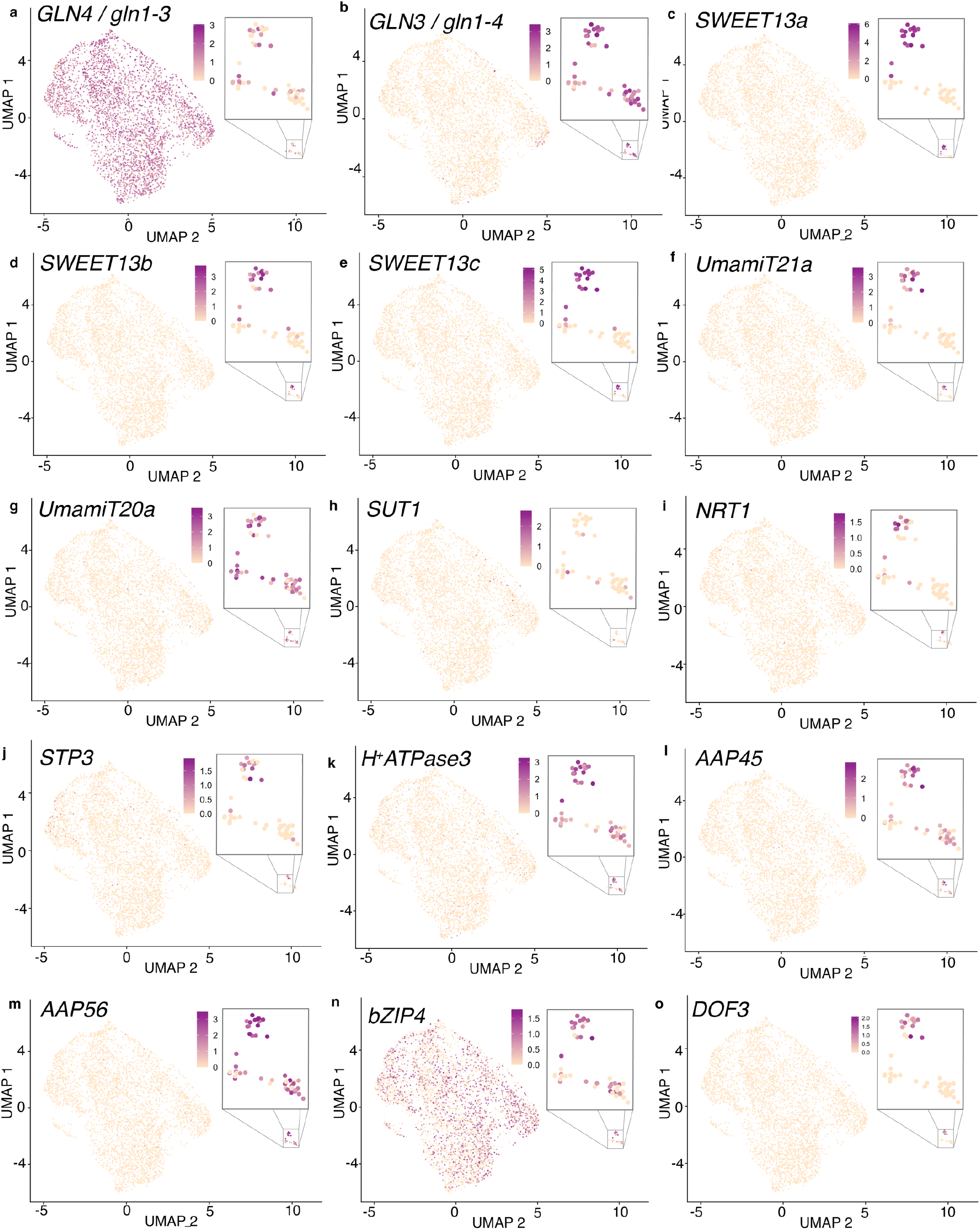
UMAP plots of glutamine synthetase, transport-related proteins, and transcription factors in bundle sheath cells. **a - b.** Feature plots show normalized levels of mRNA transcripts for glutamine synthetase genes expressed differentially in mesophyll and bundle sheath. **a.** *GLN4 (gln1-3*, protein GS3, Zm00001d017958) is widely expressed in mesophyll cells and some bundle sheath cells. **b.** *GLN3 (gln1-4*, protein GS4, Zm00001d028260) is expressed in bundle sheath cells. **c – o.** Feature plots show normalized levels of mRNA transcripts for transport-related genes and transcription factors expressed in ^ad^BS or ^ab^BS. **c.** *SWEET13a* (Zm00001d023677), **d.** *SWEET13b* (Zm00001d023673), **e.** *SWEET13c* (Zm00001d041067), and **f.** *UmamiT21a* (Zm00001d035717) are specific to ^ab^BS. **g.** *UmamiT20a* (Zm00001d044951) is expressed in ^ab^BS and ^ad^BS. **h.** *SUT1* (Zm00001d027854) is not highly expressed in any cell type in this dataset. **i.** *NRT1* (Zm00001d044768), **j.** *STP3* (Zm00001d027268), **k.** *H^+^ATPase3* (Zm00001d019062), **l.** *AAP45* (Zm00001d035243), **m.** *AAP56* (Zm00001d012231), **n.** *bZIP4* (Zm00001d018178) and **o.** *DOF3* (Zm00001d035651) are enriched in ^ab^BS relative to ^ad^BS.

**Supplementary Figure 4.**
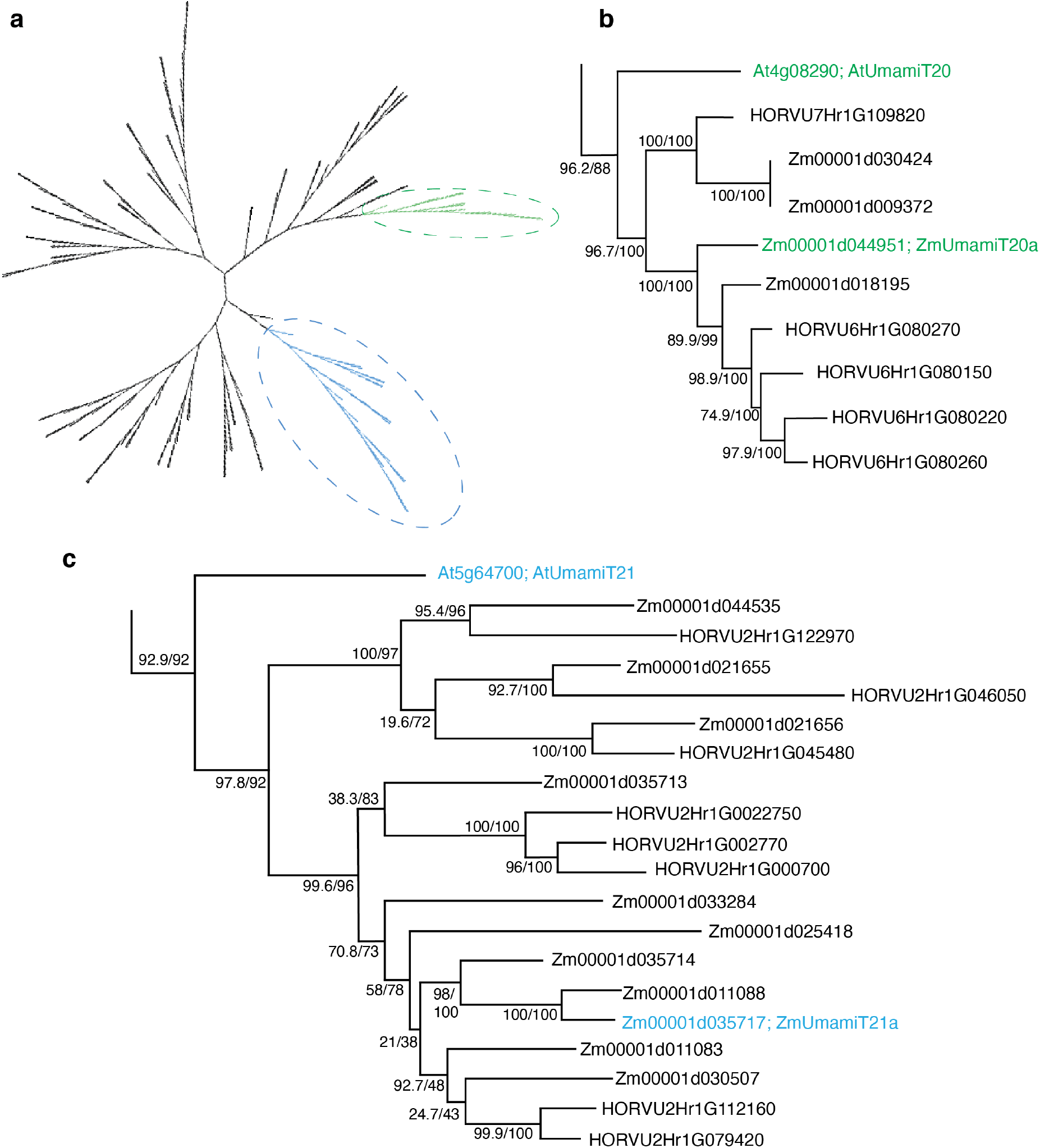
Neighbor joining tree of family of UmamiT amino acid transporters in Arabidopsis, maize, and barley. **a.** Cladogram of all UmamiT amino acid sequences in maize, barley, and Arabidopsis **b.** Phylogram of green highlighted clade containing the ZmUmamiT expressed in all BSC types. Zm00001d44951 is most closely related to Arabidopsis UmamiT20. **c.** Phylogram of blue highlighted clade containing the ZmUmamiT expressed specifically in ^ab^BSC. Zm00001d035717 is most closely related to Arabidopsis UmamiT21. Values at nodes are % UF-bootstraps out of 1000/% SH-aLRT.

**Supplementary Figure 5.**
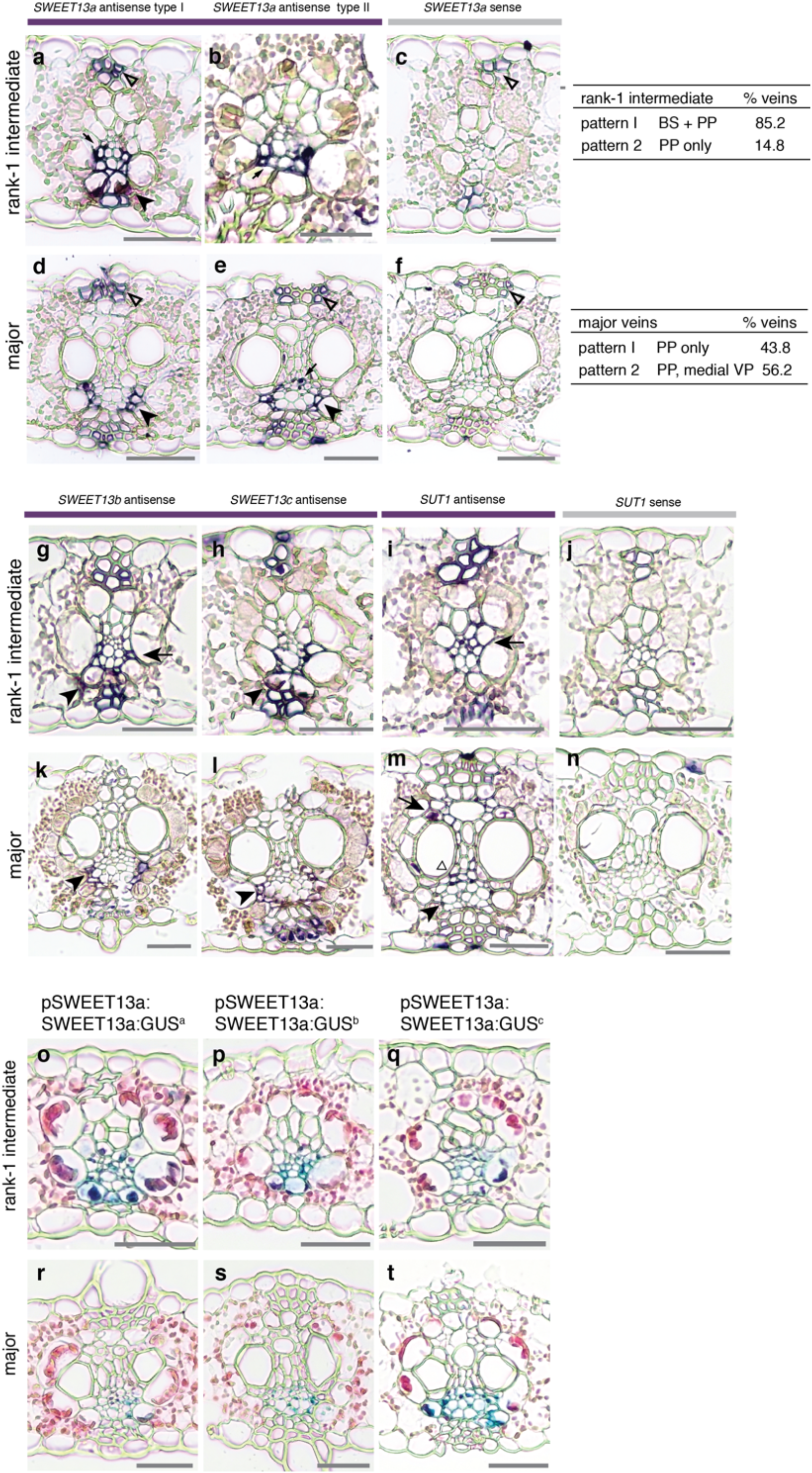
SWEET and SUT mRNA localization and SWEET13a protein localization in rank-2 intermediate and major veins. **a - c.** SWEET13a mRNA localization in rank-1 intermediate veins **a.** Pattern 1: ^ab^BSC (arrowhead) and vascular parenchyma (VP) (arrow), and **b.** Pattern 2: mainly in VP (arrow). **c**. rank-1 vein in sense-probe-hybridized section. **d-f.** SWEET13a mRNA localization in major veins **d.** Pattern 1: in phloem parenchyma (PP) only (arrow), and **e.** Pattern 2: in PP and medial VP (parenchyma between xylem and phloem) (arrow). **f.** *SWEET13a* major vein in sense-probe-hybridized section. Hypodermal sclerenchyma is considered background (open triangles). Table indicates percentage of veins meeting the above criteria for each vein type. Rank-1 intermediate, n = 196; and major veins, n = 98; variable n-numbers are due to relative proportions of each vein type in the leaf. Scale bars are 100 μm. **g - j.** *SWEET13b*, *c*, and *SUT1* mRNA localization in rank-1 intermediate veins. **g.** For *SWEET13b* antisense probe-hybridized rank-1 intermediate veins, staining was in ^ab^BS (arrowhead) and vascular parenchyma (arrow). **h.** For *SWEET13c* antisense probe-hybridized rank-1 intermediate veins, staining was in ^ab^BS (arrowhead) and vascular parenchyma (arrow). **i.** For *SUT1* antisense probe-hybridized rank-1 intermediate veins, staining was in vasculature (arrow). **j.** *SUT1* sense probe-hybridized rank-1 intermediate vein. **k - n.** *SWEET13b*, *c*, and *SUT1* mRNA localization in major veins. **k.** For *SWEET13b* antisense probe-hybridized major veins, staining was in phloem parenchyma (arrowhead). **l.** For *SWEET13c* antisense probe-hybridized major veins, staining was in phloem parenchyma (arrowhead). **m.** For *SUT1* antisense probe-hybridized major veins, staining was in companion cells (arrow), in vascular parenchyma between xylem and phloem (triangle), and in xylem parenchyma (arrowhead). **n.** *SUT1* sense probe-hybridized major vein. Scale bars are 100 μm. **o – t.** SWEET13a protein localization as visualized by GUS staining of SWEET13a:GUS-transformed B104 plants. Chloro bromoindigo precipitate is localized to abaxial portion of veins in both rank-1 intermediate (**o – q**) and major veins (**r – t**) of all three independent transformation events. Scale bars are 100 μm; sections are counterstained with Eosin-Y.

**Supplementary Figure 6.**
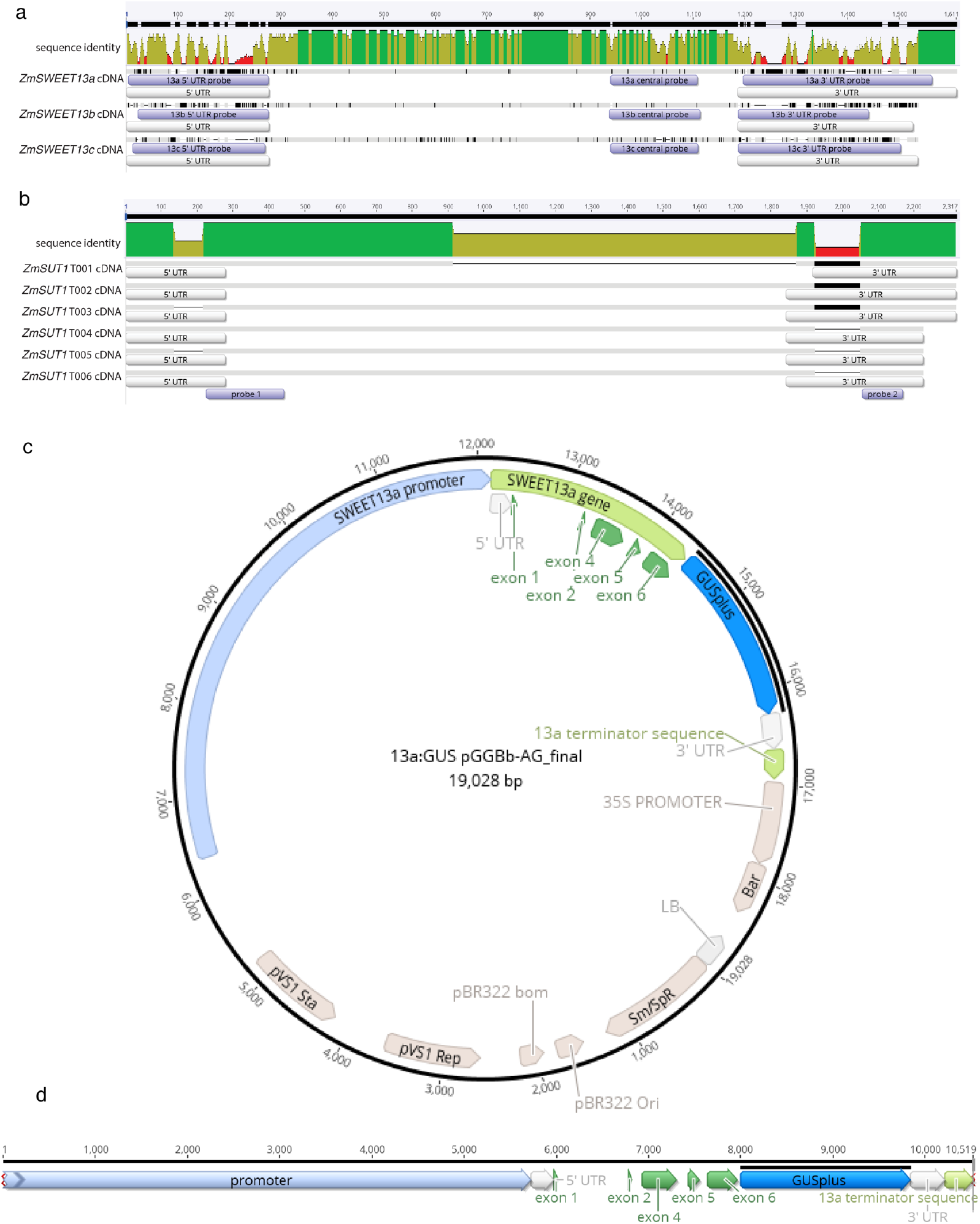
Probe design for in situ hybridization and pSWEET13a:SWEET13a:GUS construct schematic. **a.** *ZmSWEET13a*, *b*, and *c* aligned using MUSCLE in Geneious. Regions used as templates for RNA probes are highlighted in purple. Three probes in unique regions of each gene allowed us to differentiate between the homologs. **b.** The six isoforms of *ZmSUT1* aligned using MUSCLE in Geneious; two regions common to all six isoforms were selected for probe templates. Two unique regions in ZmUMAMIT21a and ZmME1 were selected for probe templates.**c.** Circular schematic of plasmid used to transform B104. The construct included the 5751 bp upstream of the start codon (lavender), all exons and introns of the SWEET13a gene (green), a 9-alanine linker fused to GUSplus (blue), followed by 684 bp downstream of stop codon (green). d. linear schematic of the SWEET13a:GUS construct

**Supplementary Figure 7.**
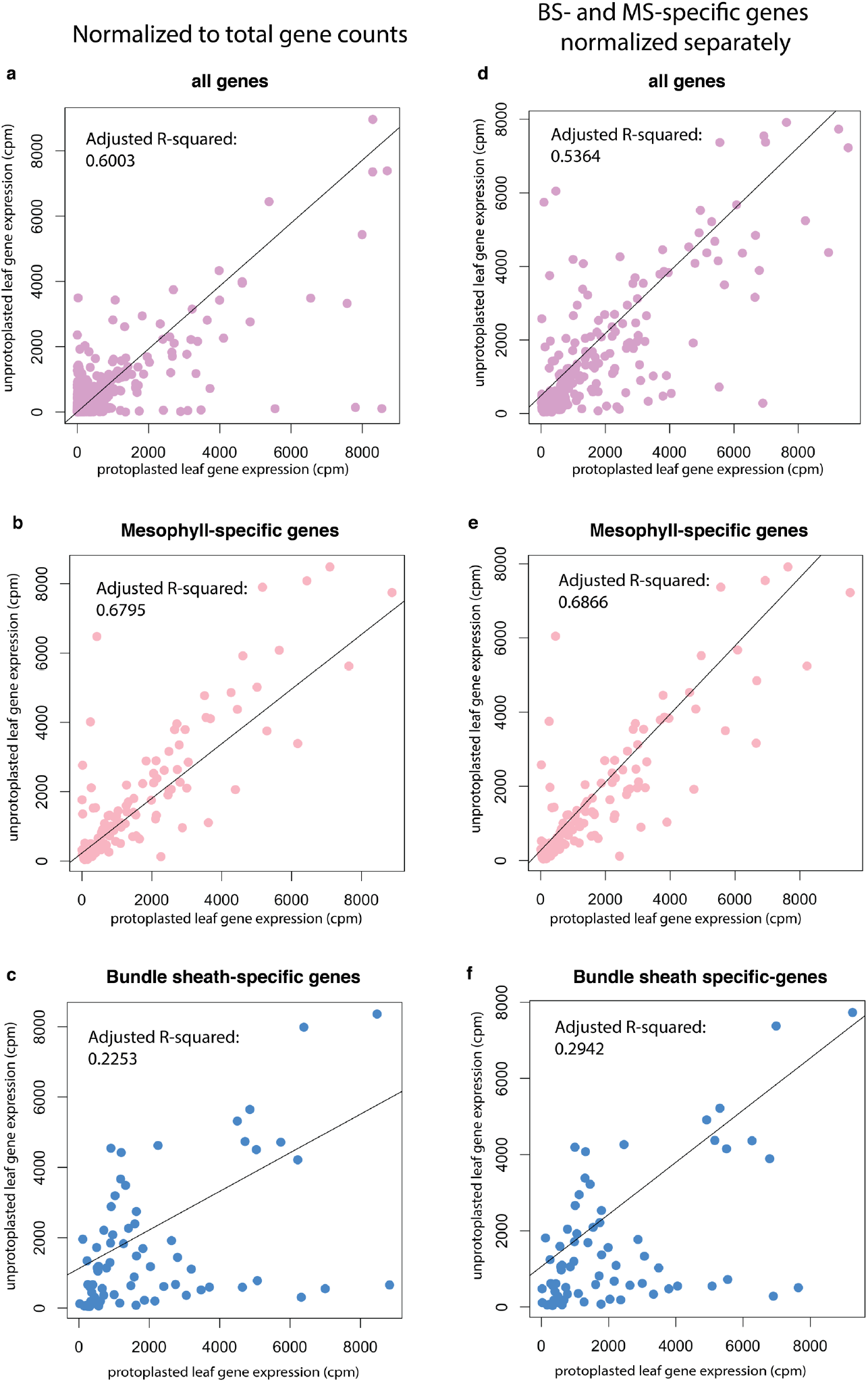
Correlation of mRNA counts between protoplasted cells and whole leaf. **a.** mRNA counts in whole leaf and protoplasted leaf expressed as counts per million reads (cpm) with adjusted R-squared values. **b**. mRNA counts for mesophyll cell-specific genes, as determined by unsupervised marker gene discovery (see materials and methods). **c.** mRNA counts for bundle sheath-specific genes. In **d-f**: counts of MS- and BS-specific genes were normalized separately to compensate for differing ratios of cell types represented in bulk leaf and protoplast samples. **d.** MS- and BS-specific genes, normalized separately. **e.** MS-specific genes normalized to total counts of MS-specific genes. **f.** BS-specific genes normalized to total counts of BS-specific genes. TMM method of normalization.

**Supplementary Table 2.**
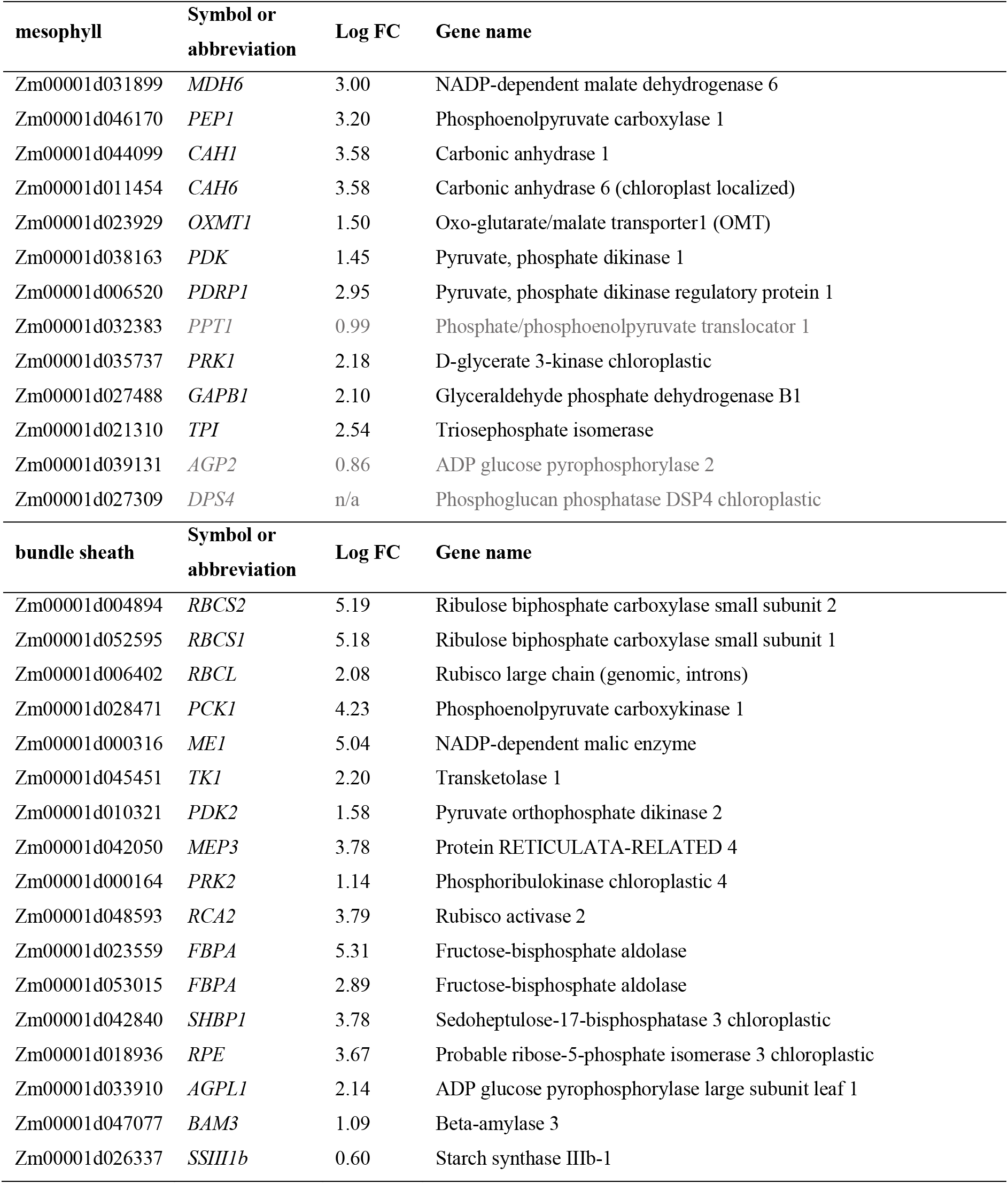
mRNA enrichment of C4 photosynthesis-related genes in MS and BS clusters.

**Supplementary Table 4.** Genes used in this study.

| **AGPv4 ID** | **gene** | **full name** |
| --- | --- | --- |
| Zm00001d023677 | SWEET13a | SWEET13a |
| Zm00001d023673 | SWEET3b | SWEET13b |
| Zm00001d041067 | SWEET13c | SWEET13c |
| Zm00001d027854 | SUT1 | Sucrose Transporter 1 |
| Zm00001d048611 | MT1b | Metallothionein-like protein 1B |
| Zm00001d038558 | CC3 | Cystatin3 |
| Zm00001d004894 | RBCS2 | ribulose biphosphate carboxylase small subunit2 |
| Zm00001d052595 | RBCS1 | ribulose biphosphate carboxylase small subunit1 |
| Zm00001d000316 | ME1 | NADP-dependent malic enzyme |
| Zm00001d053281 | NAAT1 | Nicotianamine aminotransferase |
| Zm00001d016441 | TAAT | Tyrosine aminotransferase |
| Zm00001d044099 | CAH1 | Carbonic anhydrase 1 |
| Zm00001d031899 | MD6 | NADP-dependent malate dehydrogenase 6 |
| Zm00001d046170 | PEP1 | Phosphoenolpyruvate carboxylase 1 |
| Zm00001d028471 | PCK1 | Phosphoenolpyruvate carboxykinase 1 |
| Zm00001d047658 | NAS10 | Nicotianamine synthase10 |
| Zm00001d047651 | NAS1 | Nicotianamine synthase 1 |
| Zm00001d028887 | NAS9 | Nicotianamine synthase 9 |
| Zm00001d028888 | NAS2 | Nicotianamine synthase2 |
| Zm00001d017429 | YS1 | Iron-phytosiderophore transporter yellow stripe 1 |
| Zm00001d033496 | Nas3 | Nicotianamine synthase 3 |
| Zm00001d035243 | AAAP45 | Amino acid permease 45 |
| Zm00001d012231 | AAAP56 | Amino acid permease 56 |
| Zm00001d035717 | UmamiT21a | UmamiT21a |
| Zm00001d044951 | UmamiT20a | UmamiT20a |
| Zm00001d010321 | PDK2 | Pyruvate orthophosphate dikinase 2 |
| Zm00001d038163 | PDK1 | Pyruvate orthophosphate dikinase1 |
| Zm00001d031899 | MDH6 | NADP-dependent malate dehydrogenase 6 |
| Zm00001d017958 | GLN4 | glutamine synthetase 4 (gln1-3) |
| Zm00001d033747 | GLN2 | glutamine synthetase 2 / Glutamine synthetase root isozyme 2 |
| Zm00001d026501 | GLN1 | glutamine synthetase 1 leaf isozyme chloroplastic |
| Zm00001d028260 | GLN3 | glutamine synthetase 3 / 6 / root isozyme 5 (gln1-4) |
| Zm00001d044768 | NRT1 | Protein NRT1/ PTR FAMILY 5.8 |
| Zm00001d027268 | STP3 | Sugar transport protein 3 |
| Zm00001d019062 | H<sup>+</sup>-ATPase | membrane H(+)-ATPase3 |
| Zm00001d018178 | bZIP4 | basic leucine zipper 4 / ABA-insensitive 5-like protein |
| Zm00001d010201 | MYB25 | Transcription repressor MYB6 / myb25 |

